# The representation of bodies in high level visual, prefrontal and inferior parietal cortex varies with explicit vs. implicit expression perception

**DOI:** 10.1101/2020.07.14.202515

**Authors:** Giuseppe Marrazzo, Maarten J. Vaessen, Beatrice de Gelder

## Abstract

Recent studies provide an increasingly understanding of how visual objects categories like faces or bodies are represented in the brain but also raised the question whether a category based or more network inspired models are more powerful. Two important and so far sidestepped issues in this debate concern, first, how major category attributes like the emotional expression directly influence category representation and second, whether category and attribute representation are sensitive to task demands. This study investigated the impact of a crucial category attribute like emotional expression on category activity and whether this varies with the participants’ task. Using (fMRI) we measured BOLD responses while participants viewed whole body expressions and performed either an explicit (emotion) or an implicit (shape) recognition task. Our results based on multivariate methods show that the type of task is the strongest determinant of brain activity and can be decoded in EBA, VLPFC and IPL. Brain activity was higher for the explicit task condition in VLPFC and was not emotion specific. This pattern suggests that during explicit recognition of the body expression, body category representation may be strengthened, and emotion and action related activity suppressed. Taken together these results stress the importance of the task and of the role of category attributes for understanding the functional organization of high level visual cortex.

## Introduction

Understanding how the brain processes emotion expression when these are either consciously recognized (as in standard experimental settings) or only processed implicitly (as in ongoing natural interactions) is highly relevant for assessing how body expressions influence the behavior of the observer. Category based models assume that stimulus categorization is a core process (Van Essen and Maunsell 1983; Josephs and Konkle 2020) that is relatively stable, independent from the actual task (e.g., detection, object and/or attribute identification, passive viewing, explicit recognition) and from specific stimulus attributes (e.g., emotion, gender) (Kanwisher 2017; Peelen et al. 2007). For over a decade, studies on body perception have implicitly assumed that body category representation is relatively stable and context independent and that it constitutes the gateway for processing various body attributes, similar to what has been long argued for face categories (Shallice 1988; Kanwisher 2000; Kanwisher and Yovel 2006; Kanwisher, McDermott, and Chun 1997; Peelen and Downing 2007; de Gelder and Poyo Solanas 2021). Available evidence shows that body expression perception is associated with activity in ventral body areas as well as in areas outside the body selective ones. (de Gelder 2006; Goldberg, Preminger, and Malach 2014)

On the other hand, a less category-centric picture may be more suited for addressing task variable and for understanding how category attributes are processed. There is growing evidence showing that task settings significantly impact the activity in object category areas, including body selective ones. For example, selective attention-related increases have been found in category representation areas for the preferred category during visual search tasks. (Cukur et al. 2013; Peelen, Fei-Fei, and Kastner 2009). There is increasing evidence that the brain encodes stimulus information in high-dimensional representational spaces based on the joint activity of neural populations (Averbeck, Latham, and Pouget 2006; Haxby, Connolly, and Guntupalli 2014; Kriegeskorte et al. 2008). This encoding process may be dynamic, relatively task sensitive and at the service of different and complex behavioral goals (Hebart et al. 2018). The emerging network picture is a change from more static views of category representation favoring dedicated functional areas (Betzel 2020).

Attribute representation and task sensitivity are two important issues in this debate. First, it is currently an open question to what extent specific body attributes, like identity or emotional expression, influence the activity and selectivity of body areas in ventrotemporal cortex, extrastriate body area (EBA) and the more anterior fusiform body area (FBA) (Ross and Flack 2020; de Gelder and Poyo Solanas 2021; Peelen and Downing 2017). Studies of body expression perception have systematically reported an impact of emotional expression on activity in EBA and FBA (Peelen and Downing 2007; Pichon, de Gelder, and Grezes 2009, 2012; Hadjikhani and de Gelder 2003). Different from EBA, FBA has been suggested to have a bigger involvement in identity and emotion processing through its connections to other areas, like the amygdalae (Orgs et al. 2015). EBA and FBA may also have different roles for different emotions. For example, Peelen and colleagues found that fear significantly modulated EBA but not FBA while no difference was found in activity patterns for other expressions (Peelen et al. 2007). Such emotion specific differences have been linked to differences in attention, arousal etc. For example, it has been shown that the strength of emotion modulation in FBA is related, on a voxel-by-voxel basis, to the degree of body selectivity and is positively correlated with amygdala activation (Peelen et al. 2007). Most interestingly, the fact that EBA seems more sensitive to fearful body expressions than FBA makes more sense from a biological survival point of view defining emotions as action plans (Frijda 1986) and EBA has been suggested to be the interface between perceptual and motor processes (Orgs et al. 2015).

Second, it is still poorly understood whether expression sensitivity of the body areas itself varies with the task, ie. whether the specific task changes how a body area represents the emotion of the body stimulus. It has been argued that the task impacts processing in prefrontal and parietal areas but not necessarily in ventral temporal category selective areas (Bugatus, Weiner, and Grill-Spector 2017; Tsotsos 2011; Bracci, Daniels, and Op de Beeck 2017; Xu and Vaziri-Pashkam 2019). More specifically, the task may require explicit recognition of a body attribute like the emotional expressions as opposed to incidental or implicit perception where no recognition of the expression is asked for. A classic example of implicit processing task is a gender recognition task used for measuring implicit processing of facial expressions (e.g. (Vuilleumier et al. 2005) or a color monitoring task used or implicit perception of body expressions (Pichon, de Gelder, and Grezes 2012). For instance, we observed increased activity in FBA and EBA when participants performed an emotion versus a color-naming tasks with whole body videos (Pichon, de Gelder, and Grezes 2012; Sinke et al. 2012). Implicit processing is also related to exogenous attention or stimulus driven attention, a well know source of representational dynamics (Carretie 2014). Affective stimulus attributes modulate the role of attention as shown for example with findings that bodies with fear expressions have different effects on saccades than neutral bodies (Bannerman et al. 2009) and that in hemispatial neglect patients, contralesional presentation of fear body expressions reduces neglect (Tamietto et al. 2015). In an effort to disentangle the effects of attention and task, (Bugatus, Weiner, and Grill-Spector 2017) showed that attention has an influence on category representation in high level visual cortex and in prefrontal cortex, while task did influence activity in prefrontal cortex but not in high level visual cortex. As concerns stimulus awareness, activity in ventral body category representation areas is significantly reduced for unaware stimuli but remains the same in dorsal action representation areas (Zhan, Goebel, and de Gelder 2018).

The goal of this study was to investigate whether the type of task and of emotion expression influences the representation of bodies and body expressions inside and outside body selective category areas during measurement of brain activity with fMRI. We used decoding analysis to discover how body areas are involved in explicit as opposed to implicit expression processing. If ventrotemporal body object categories areas (EBA, FBA) are relatively insensitive to task dynamics then they should not be among the areas where task differences are observed. Alternatively, body category representation areas may be directly involved in expression recognition or indirectly through their connectivity with other important brain areas that are known to play a role in expression processing like the amygdalae (Vuilleumier et al. 2004; de Gelder, Hortensius, and Tamietto 2012), prefrontal areas (VLPFC) and action representation areas in parietal cortex, specifically intraparietal sulcus (IPS) and inferior parietal lobule (IPL).

Two different tasks were designed to be formally similar (similar difficulty, similar response alternatives) for use with the same stimulus materials that consisted of body expressions with two different emotions and two different skin colors. One task, emotion perception, required explicit recognition of the body expression and a forced choice between two alternatives. The other task was shape perception and required explicit recognition of a shape overlaid on the body image and a forced choice between two shape alternatives. We used multivariate decoding and RSA in order to decode stimulus and task related information in locally defined patterns of brain activity (Connolly et al. 2012; Connolly et al. 2016; Huth et al. 2012; Kriegeskorte et al. 2008; Mitchell et al. 2008; Nastase et al. 2017; Oosterhof et al. 2010; Sha et al. 2015). Our goal was to answer the question whether activity in body category representation areas EBA and FBA would vary significantly between the emotion vs the shape task and whether this difference could also be decoded in other areas possibly in amygdalae. The alternatively outcome would be that the task cannot be decoded in the category areas, indicating that category representation is immune from task requirements and attribute recognition. To anticipate, our results show that the difference between the two tasks can be decoded in EBA, VLPFC and IPL and that task sensitivity but not attribute selectivity is clearly seen in category selective areas in the higher visual cortex and in the VLPFC.

## Materials and Methods

The present study uses brain and behavioral data previously collected and described in (Watson and de Gelder 2017) but now analyzed from a different theoretical perspective and with fully different methods.

### Participants

Data of twenty Caucasian participants were used for the current study (8 males, mean age ± standard deviation=22 ± 3.51 years). Participants were naive to the task and the stimuli and received a monetary reward for their participation. Written informed consent was provided before starting the protocol. The scanning session took place at the neuroimaging facility Scannexus at Maastricht University. All procedures conformed with the Declaration of Helsinki and the study was approved by the Ethics Committee of Maastricht University.

### Stimuli

Stimuli consisted of still images of angry and happy body postures of black African and white Caucasian ethnicity. The set of black body expressions was obtained by instructing black African participants, all residents of Cape Town, South Africa, to imagine a range of daily events and show how they would react to them nonverbally. The set of white affective body stimuli (five males each expressing anger and happiness) were selected from a set previously validated (Stienen, Tanaka, and de Gelder 2011; Van den Stock et al. 2011). Both sets were pre-processed with the same software and underwent the same post-selection procedure. Photographs were captured using a Nikon V1 35mm camera equipped with a Nikon 30-100mm lens on a tripod, and under studio lighting. The stimulus set consisted of 20 affective bodies (2 races (Black, White) x 2 emotions (Angry, Happy) x 5 identities). The photos showed the entire body, including the hands and feet. For behavioral validation of the images ten white European participants were then asked to categorize the emotion expressed in a given picture (neutrality, anger, happiness, fear, sadness, disgust). All emotions were recognized above 70%. Based on these results five male identities were chosen, with photos of the same identity expressing both anger and happiness. Ten upright white and black (20 in total) affective body images were selected for the final stimulus set. Pictures were edited using Adobe Photoshop CC 14 software (Adobe System Incorporated) in order to blur the faces using an averaged skin color; thus, there was no information in the face.

### fMRI Acquisition and Experimental Procedure

Participants were scanned using a Siemens 3T Prisma scanner. Padding and earplugs were used to reduce head movements and scanner noise. Stimuli were projected to the center of a semi-translucent screen at the back of the scanner bore that participants could see using a mirror mounted on the head coil. Participants were instructed to fixate on the geometrical figure overlaid on the stimulus which was positioned on the most neutral or least informative part of the body. Given this arrangement no extra fixation cross was added on top of the geometrical figure. In between trials the fixation cross was present and in experimental trials the geometric figure served as fixation point.

The experiment comprised two tasks presented in a mixed block/event related design of four separate runs. Each run consisted of a presentation of emotion (A) and shape (B) blocks (AB – BA – BA – AB) and in each block stimuli were presented in a slow event related manner. The two different tasks were designed to provide information on explicit and implicit emotion perception. For the emotion block, participants were instructed to respond on whether the emotion expressed was anger or happiness. In the shape block, participants judged whether the stimulus contained a circle or a square which was superimposed on the body. The task was indicated on the screen for 2 s before each block began. The trials in each block were separated by a fixation cross on a gray background that appeared for 10 or 12 s (in a pseudo-random order). Following the fixation cross, a body image was presented for 500 ms (during each trial the participants were instructed to fixate) followed by a response screen lasting 1500 ms, showing the two response options on the left and right of the fixation cross and corresponding to the index and to the middle finger respectively. The side of the response options were randomized per trial to avoid motor preparation. Each stimulus was presented twice in each run, once during the emotion task and once during the shape task. Thus, each run consisted of 40 trials (+ 2 task indicators), see Fig. 1.

**Figure 1.**
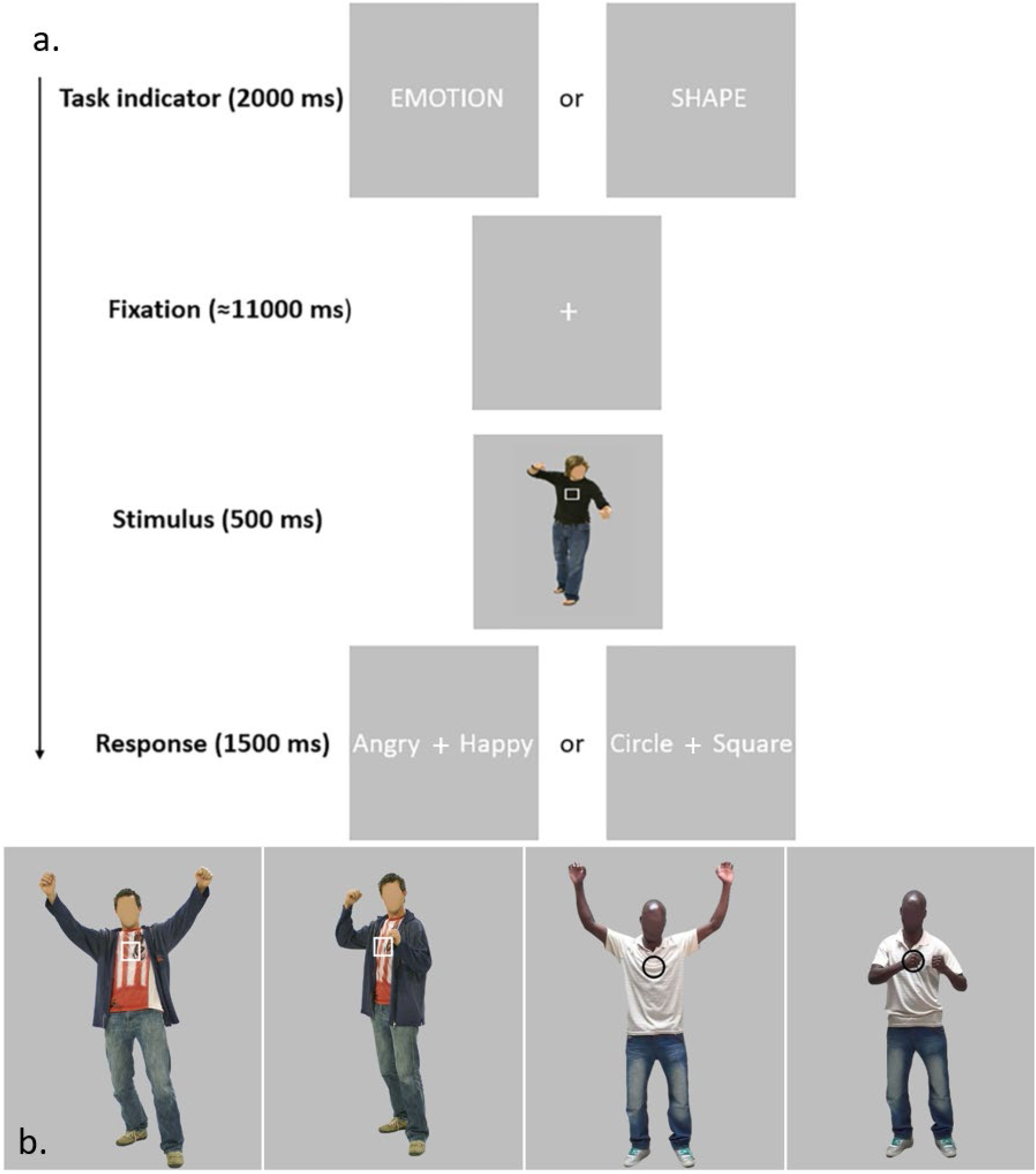
**(a) Examples of explicit and implicit trials.** During the experiment a task indicator appeared (2000 ms) showing which task (explicit emotional evaluation or implicit emotional evaluation) the participants were asked to perform. The task indicator was followed by a fixation period, the stimulus (white happy/angry, or black happy/angry) and a response window. Participants responded via two buttons pressed by the index finger (word on the left) and the middle finger (word on the right), with randomization of the response options in order to avoid motor preparation (*Watson and de Gelder 2017*). **(b) Example of different angry (happy) poses.** Four different examples of unique affective body poses depicting happiness (first picture and third picture from the left) and anger (second picture and forth picture from the left). Participants were asked to recognize the emotion in the explicit task and name the shape (square/circle superimposed) in the implicit task.

### MRI acquisition and Data Preprocessing

A T2*-weighted gradient echo EPI sequence was used to acquire the functional data covering the whole brain with 2 x 2 x 2 mm^3^ resolution (64 slices without gaps, TR = 2000 ms, TE= 30 ms, flip angle= 77 °, multiband acceleration factor = 2, FOV = 160 x 160 mm, matrix size = 100 x 100). Furthermore, a T1-weighted MPRAGE sequence was used for each participant (1 x 1 x 1 mm^3^, TR=2300 ms, TE= 2.98 ms). Preprocessing was performed using BrainVoyager software (BrainVoyager QX) (Brain Innovation B.V., Maastricht, the Netherlands). For each run a slice scan time correction using sinc interpolation was performed, data from each run was motion-corrected by realigning to the first volume of the first run using sinc interpolation. A two-cycle temporal high-pass filtering was applied in order to remove low frequency linear and quadratic trends. Notice that no spatial smoothing was performed at this stage. The anatomical data, after the skull removal and inhomogeneity correction, were spatially warped to MNI space (MNI-ICBM 152), and the functional data were then co-registered to the anatomical data in the new space using the boundary based registration algorithm (Greve and Fischl 2009).

### Univariate Analysis

Using BrainVoyager (BV, v21.2) we first defined a subject-specific univariate general linear model (GLM) where each condition (emotion black angry (E_BA), emotion black happy (E_BH), emotion white angry (E_WA), emotion white happy (E_WH), shape black angry (S_BA), shape black happy (S_BH), shape white angry (S_WA), shape white happy (S_WH)) was included as a square wave of the same duration of the trial, convolved with the canonical hemodynamic response function. The 3D motion parameter estimates were included as regressors of no interest in the design matrix. For the group statistical analysis, we first performed spatial smoothing with a Gaussian Kernel (3 mm) of all the functional images and then, in order to assess the variability of observed effects across subjects, we combined the individual GLM’s in a random effects (RFX) GLM analysis, as is the custom in the BV pipeline. For 7 participants, only three of the five original trials for each condition were included as predictors due to an initial error in stimulus presentation, resulting in a reduced set of 96 trials out of 160 (2 emotions x 2 skin color x 2 tasks x 5 repetitions x 4 runs). To test for effects and interactions between the factors an RFX three-way repeated measures ANOVA was performed in BV on the combined individual GLM’s.

### Multivariate Analysis

All multivariate analyses were conducted with in-house MATLAB scripts (vR2018a, The MathWorks Inc., Natick, MA, USA). First, the BOLD time course of each voxel was divided in single trials, whose temporal window (epoch) were defined between 1TR prior and 4TR after the stimulus onset, resulting in 42 trials per run (168 in total). Within each run, 2 trials represented the task indicator and therefore they were not included in the analysis. Each trial was normalized with respect to the baseline 2000 ms, before the first stimulus onset (the first TR in the trial segment). We linearly fitted the percent BOLD signal change of each voxel and each trial separately with a design matrix consisting of a constant term (intercept) and an optimized hemodynamic response function (HRF). The optimized HRF was designed to take into account potential differences in the BOLD responses (temporal delay) for a certain voxel. The optimal delay was calculated for each voxel by convolving a canonical HRF with a box-car predictor whose value was one when the stimulus was presented. The time-to-peak parameter was varied between 4.0 s and 6.0 s in steps of 0.5 s. The five resulting HRFs were fit to the percent BOLD signal change of all trials averaged and the time-to-peak giving the best fit was chosen as the optimal HRF delay of that voxel. For each trial and each voxel, we then used the resulting β-values as a feature in the classifier (Gardumi et al. 2016). The method provided above does not represent the standard procedure for multivariate analysis in which β-values from the univariate GLM are used as feature in the classifier. The traditional GLM uses a fixed parameter modelling the positive time to peak of the HRF and the estimated β of the responses to each category are used for statistical inference. Although the statistical framework is not available for the optimized HRF method, the multivariate classifier can work both with the traditional GLM β and the HRF optimized β. Furthermore, the optimized HRF method has clear advantage compared to the standard framework, because it estimates with higher precision the delay of the canonical HRF used to model the response (5 possible choices within the standard range of variation of the positive time to peak: 4.0 – 6.0 s).

#### Searchlight analysis

In order to perform whole brain decoding (Kriegeskorte, Goebel, and Bandettini 2006) we implemented the method proposed by (Ontivero-Ortega et al. 2017), in which the brain is divided into spheres of searchlights and a fast Gaussian Naïve Bayes (GNB) classifier is fitted in each of them. Each searchlight has a radius of 5 voxels and is defined by a central voxel and a set of voxels in its neighborhood. The classification accuracy of the searchlight region was then assigned to the central voxel. In order to avoid overfitting, for each subject we split the data following the leave-one-run-out paradigm (4 – fold cross-validation) and computed the prediction accuracy by testing the trained classifier on left-out test data. The GNB classifier was trained to predict tasks (Emotion vs Shape), emotion (Angry bodies vs Happy bodies) or skin color (Black bodies vs White bodies). Here the responses to individual stimuli were averaged for the 8 main conditions of the experiment. The emotion and skin color effects decoding were determined both across the tasks (160 trials available for training and testing the classifier) and within the tasks (80 trials for the explicit task, 80 trials for the implicit task), for 7 participants (see Univariate analysis) only 96 trials out 160 were available for the analysis. Moreover, in order to determine interstimulus differences in the multivoxel patterns (MVPs), the GNB was trained to classify the 20 unique affective bodies (5 identities x 2 skin colors x 2 emotions).

#### Interstimulus decoding

In order to check whether the qualitative differences in the 20 unique poses (5 identities x 2 skin color x 2 emotions) of the stimulus set were also reflected in the MVPs, a GNB classifier was trained to classify the 20 affective bodies. Specifically, for each searchlight we assigned a unique label to each different stimulus and trained the GNB to classify it following the leave-one-run-out paradigm. We then assessed the ability of the classifier to categorize the different poses on the left-out data, by assigning the corresponding prediction accuracy value to the central voxel of each searchlight.

#### Whole brain RSA of intra-versus inter-conditions similarities analysis

In addition to decoding with a classifier, another method to detect condition effects in MVP’s is to statistically test for differences between intra-versus inter-condition MPV similarities (Peelen, Atkinson, and Vuilleumier 2010). As in the GNB analysis, for each subject and for each 5 voxels radius searchlight spanning the whole brain, we built neural representational dissimilarity matrices (RDMs) by computing the dissimilarity (1 - Pearson’s correlation) between the multivoxel patterns of each of the 160 trials. Next, we extracted from these RDMs the intra-condition or inter-condition elements and compared these with a two-sample t-test. This test was performed for the conditions of task, emotion and skin color separately. Furthermore, we assessed task specific differences between intra-versus inter-condition MVP similarities by extracting neural RDMs for emotion and skin condition within the explicit and implicit task separately. This was performed by testing the task specific neural RDMs (80 trials per task). As mentioned in the univariate analysis, for 7 participants 2 trials for each condition were to be discarded, resulting in 96 trials (48 per each task). On a group level, for each voxel, single-subject results were tested against zero, resulting in a group two-tailed t-test.

### Group Analysis

For the group-level analysis spatial smoothing (Gaussian kernel of 3mm FWHM) was applied to the resulting maps of each individual. For the decoding analysis with the GNB classifiers the maps contained the classification accuracies minus chance level and for the inter-versus intra-condition MVP similarity analysis the maps represented the t-values from the t-test. Next, for all analyses, a statistical map was obtained by performing a two tailed t-test against zero over subjects. The statistical threshold for the overall activation pattern was q = .05 corrected for multiple comparison using the false discovery rate (FDR).

### Region of Interest Analysis

Regions of interest (ROIs) were defined in a 5-fold cross-validation procedure performed as follows. For each fold, single subject accuracy maps produced by GNB decoding on task effect were split in two sets: a training set of n=16 and test set of n=4 respectively. The larger n=16 set was used for defining the ROIs and the smaller n=4 set for extracting MVPs. Data from each training set were tested on the group level against chance level of accuracy in a t-test, the resulting t-map was thresholded in BrainVoyager at q(FDR) = .01 and the coordinates of each peak voxel cluster were extracted (see Table S8 in supplementary material). This statistical threshold allowed us to obtain spatially separated clusters across each fold from which we extracted the peak voxel coordinates. We defined a sphere of radius r = 8 voxels around each peak value and all the voxels within the sphere whose t-value was above the threshold of q(FDR) = .05 were selected as part of the ROI (see Fig. 8a). Subsequently, multivoxel patterns from the ROIs defined above were extracted from the testing set (4 left-out subjects). We computed Representational Dissimilarity Matrices (RDMs) via a metric of distance (1 – Pearson’s correlation coefficient) between the multivoxel patterns of the left-out subjects from the 8 conditions of the main experiment. Additionally, for each ROI and for each fold, to assess the overall activation level we plotted the beta values from the optimized HRF model for the different experimental conditions. We extracted beta values from the left-out subjects within each ROI (see above) by averaging the multi voxel patterns of each condition. Within each fold the 4 sets (one for each of the subject in the test set) of RDMs and beta values were the again averaged. We repeated the procedure described above 5 times permuting the subjects belonging to the training set (define ROI) and the test set (extracting responses). Ultimately, beta values and RDMs were averaged across the 5 folds resulting in 2 plots for each ROI (see Fig. 8b).

## Results

### Behavioral analysis

To test for any difference in performance between the two emotion and shape tasks we performed a three-way repeated measure ANOVA on accuracies and response times completing the previous results (Watson and de Gelder 2017). For each subject we averaged the 8 conditions over repetitions. The analysis on the accuracies revealed a main effect of the three factors task, skin and emotions (F(1,19) = 40.06, p < .001; F(1,19) = 28.88, p < .001; F(1,19) = 14.08, p = .001). In order to check the direction of the effect, a paired sample t-test was performed. The latter revealed that the mean accuracy for the emotion task was significantly smaller than the mean accuracy for the shape task (mean emotion = .893 ± .156, mean shape = .986 ± .027, t(79) = −5.050 p < .001). Likewise, we found that the mean accuracies for the angry poses and the black poses (averaged across the tasks) were significantly lower than the mean accuracies for the happy poses and the white poses respectively (mean angry = .910 ± .158, mean happy = .969 ± .052, t(79) = −3.243 p = .002; mean black = .911 ± .155, mean white = .968 ± .063, t(79) = −2.904 p = .005). The complete results are reported in the supplementary material (see Table S2, S4, S5).

The analysis on the response times showed a main effect of task and emotion (F(1,19) = 34.58, p < .001; F(1,19) = 6.76, p = .018). A paired sample t-test revealed that the mean response time for the emotion task was significantly greater compared to the shape task (mean emotion = 843.01 ± 111.77 ms, mean shape = 717.35 ± 85.44 ms, t(79) = 8.63 p < .001) and the mean response time for the angry was significantly higher than the happy conditions (mean angry = 796.61± 130.25 ms, mean happy = 763.75 ± 101.37 ms, t(79) = 2.94, p = .004). Furthermore, task affects the response times for the emotion conditions and for the skin conditions (F(1,19) = 4.66, p = .044; F(1,19) = 30.33, p < .001). When participants explicitly named the emotion, we found a significant difference in the response times with more time needed to name an angry compared to a happy image (mean angry = 873.65 ± 114.80 ms, mean happy = 812.37 ± 101.01 ms, t(39) = 3.23, p = .002). This difference was not significant during the shape categorization task. For the emotion categorization condition response times were longer for the black stimuli (mean black = 875.30 ± 102.18ms, mean white = 810.72 ± 112.82 ms, t(39) = 4.25, p < .001). In contrast, for the shape categorization task mean response time for white conditions were longer that for the black stimuli (mean black = 706.04 ± 84.37 ms, mean white = 728.66 ± 86.06 ms, t(39) = −2.28, p = .002). The complete results are reported in the supplementary material (see Table S3, S6, S7). Taken together these behavioral results show significant differences between conditions, but the actual order of magnitude is such that, at this very high accuracy level, this difference although statistically significant does not reflect a substantial, meaningful behavioral distinction between the tasks. Moreover, these are not reaction times as a delayed naming task was used.

### Analysis of condition effects in activation level

In the univariate analysis we tested the effect of the 3 main factors (task: explicit vs implicit; emotion: angry vs. happy; skin color: black vs. white) and their interactions, and in order to determine the direction of the effect we computed a two-tailed t-test on each pairwise contrasts. We found significant higher responses for the explicit task in lateral occipito-temporal cortex (LOTC), medial superior frontal gyrus (MSFG), bilateral ventrolateral prefrontal cortex (VLPFC) and bilateral anterior insular cortex (AIC). Higher activation levels for the implicit task were found in bilateral superior temporal gyrus (STG), right middle temporal gyrus (MTG), right inferior parietal lobule (IPL), bilateral marginal sulcus (MS) and left anterior cingulate cortex (ACC) (see Fig. 2a and Table 1). The contrast Angry vs. Happy bodies for all trials as well as for the emotion task trials only, revealed higher activation for happy bodies in the primary visual cortex (MNI: - 13, −81, −9; t(19) = −8.01, p <.001) (see Fig 2b). No significant differences in activation levels were found for Black vs. White bodies. The ANOVA showed that the only interaction which gave above threshold (q(FDR)<.05) clusters was the one between emotions and skin color (table S1 in supplementary material) see also (Watson and de Gelder 2017) for the details.

**Figure 2.**
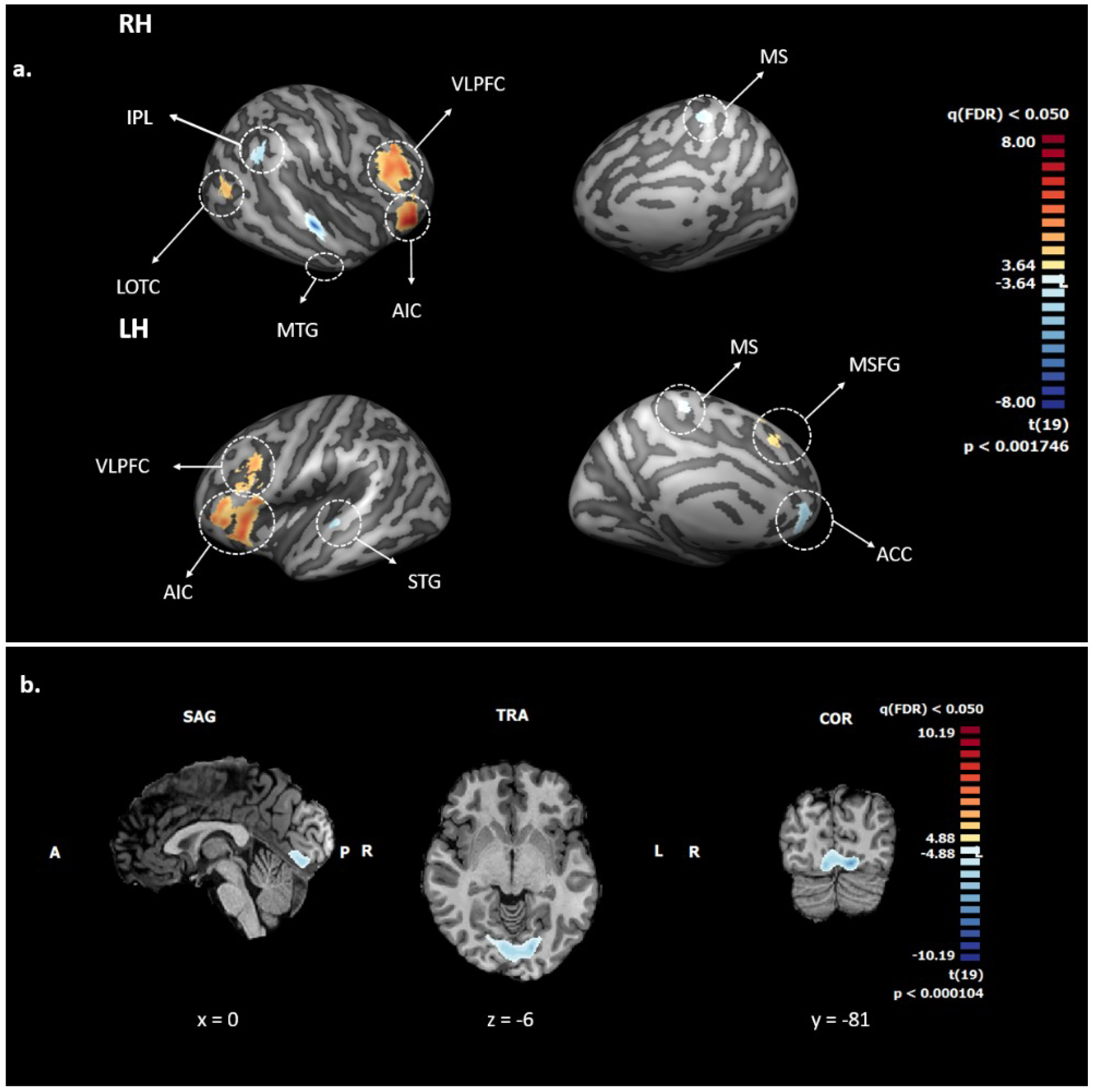
**(a): Whole Brain Analysis: Univariate results for explicit vs. implicit expression recognition task (q(FDR) < .05).** The color map indicates regions where higher (red) or lower (blue) activation was found for the emotion recognition task (explicit) vs the shape recognition task (implicit). Statistical analysis was performed on the volume maps and the resulting brain regions, after thresholding, are mapped to and overlaid on the inflated group average cortical surface for visualization purposes. Abbreviations: ACC = anterior cingulate cortex, AIC = anterior insular cortex, IPL = inferior parietal lobe, LOTC = lateral-occipitotemporal cortex, MS = marginal sulcus, MSFG = medial superior frontal gyrus, MTG = middle temporal gyrus, STG= superior temporal gyrus, VLPFC = ventrolateral prefrontal cortex. **(b): Whole Brain Analysis: Univariate results for angry vs. happy expression recognition task (q(FDR) < .05).** The color map indicates regions where higher (red) or lower (blue) activation was found for the angry body pose vs happy body pose averaged across the tasks. One cluster was found spanning the early visual area with higher activation for happy bodies.

**Table 1.** Whole Brain Group level univariate results of explicit vs. implicit conditions. The table shows the regions where greater activity was found for the explicit conditions (t>0) and the implicit conditions (t<0). The t-map was thresholded at q(FDR) < .05 and cluster size corrected. Peak voxel coordinates (MNI) and corresponding t value of each surviving cluster are reported. The degrees of freedom for the t-test were 19 while for the ANOVA 1 and 19. All the results were significant at p < .001.

| Brain Regions | L/R | x | y | z | t(19) | F(1,19) |
| --- | --- | --- | --- | --- | --- | --- |
| Superior temporal gyrus | R | 65 | -16 | 1 | 8.678*** | 75.525*** |
|  | L | -68 | -8 | -3 | -7.021*** | 45.418** |
| Middle temporal gyrus | R | 59 | -11 | -36 | -6.173** | 38.140** |
| Inferior parietal lobule | R | 47 | -47 | 32 | -5.043* | 25.471* |
| Lateral occipitotemporal cortex | R | 53 | -66 | 13 | 6.127** | 37.647** |
| Marginal sulcus | R | 6 | -30 | 54 | -5.396* | 29.219* |
| Ventrolateral prefrontal cortex | R | 45 | 25 | 18 | 8.684*** | 75.587*** |
|  | L | -45 | 17 | 25 | 5.734* | 32.934* |
| Medial superior frontal gyrus |  | 0 | 18 | 59 | 5.831* | 34.040* |
| Anterior cingulate cortex |  | 0 | 33 | -11 | -5.667* | 32.173* |
| Anterior insular cortex | R | 36 | 26 | -3 | 7.615*** | 57.663*** |
|  | L | -34 | 22 | -3 | 6.368** | 40.571** |

\* $p < .0001$
\*\* $p < .00001$
\*\*\* $p < .000001$

### Multivariate decoding of task effect

The whole brain searchlight GNB analysis revealed significant above-chance classification of the explicit vs. implicit task at the group level in bilateral lateral occipito-temporal cortex (LOTC), bilateral posterior inferior temporal gyrus (PITG), posterior middle temporal gyrus (PMTG), right inferior parietal lobule (IPL), bilateral ventrolateral prefrontal cortex (VLPFC), precuneus (PCUN), posterior cingulate cortex (PCC), fusiform gyrus (FG), medial superior frontal gyrus (MSFG) and cerebellum (CB) (See Fig. 3 and Table 2 for details). Moreover, these regions overlapped substantially with the univariate GLM results as shown in Fig. 5a. Importantly, the extent and statistical significance of the multivariate GNB results where much larger than for the GLM analysis, possibly indicating that the task effect was not only expressed through the level of activation but also in different multi-voxel patterns (regardless of level of activation). We also performed an analysis of the angry vs. happy bodies decoding (trials of both tasks combined) and found above chance classification accuracies in the right FG (MNI: 29, −49, −20; t(19) = 5.80, p < .001), and cerebellum (MNI: 29, −43, −34; t(19) = 4.90, p < .001). When considering the tasks separately, we did not find any regions where emotion could be decoded. When decoding angry vs. happy bodies (for each task separately) and black vs. white bodies (trials of both tasks combined, and for each task separately) the classification did not yield any above chance results at the group level.

**Figure 3.**
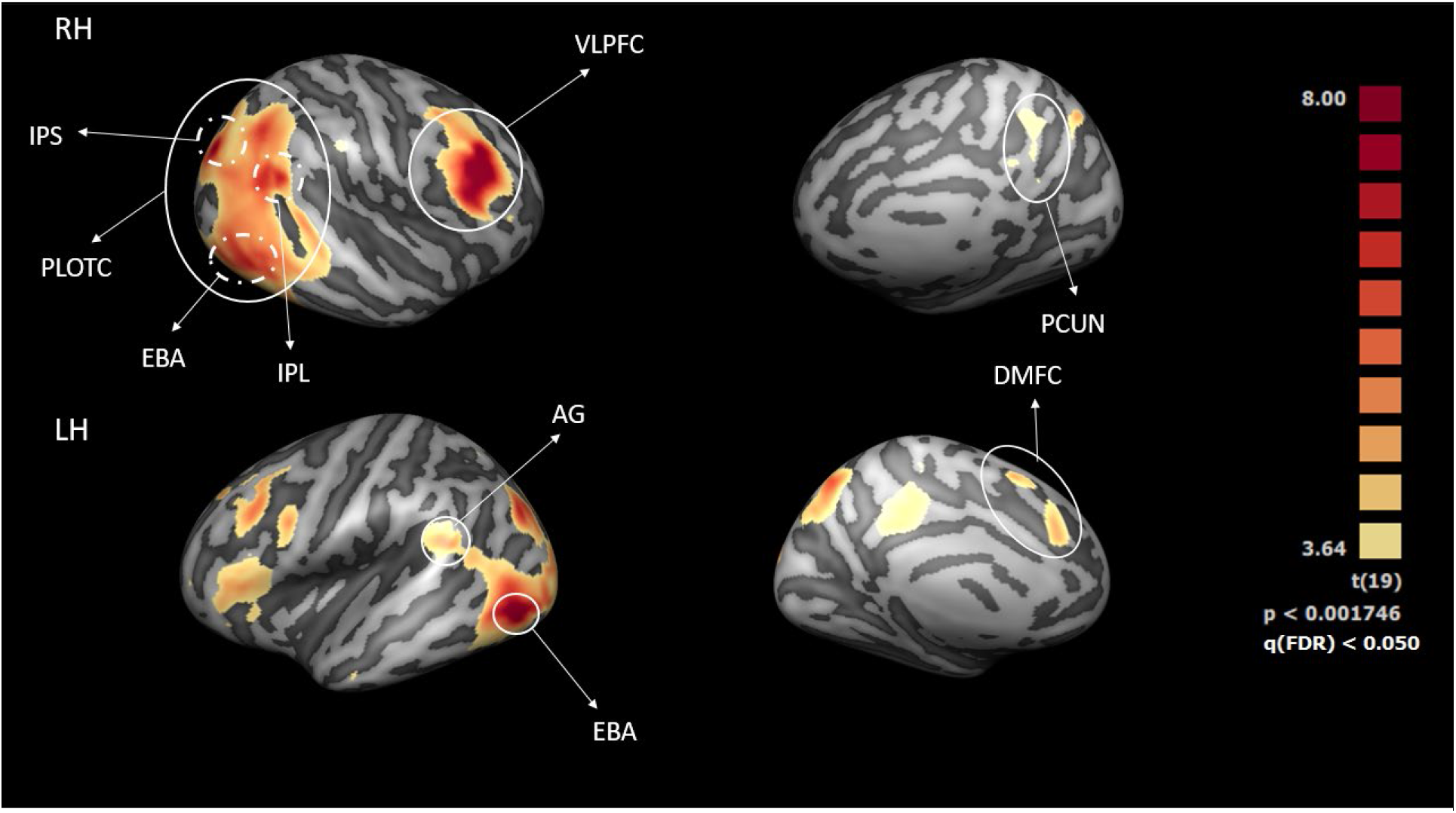
Whole Brain MVPA Analysis: results of the GNB classifier for explicit vs. implicit task. Above chance classification accuracies produced by the searchlight GNB, q(FDR) < .05 and cluster size corrected (min. cluster size threshold = 176) are shown. The color map indicates the t-value of the test against chance level accuracy. Abbreviations: AG = angular gyrus; DMFC = dorsomedial frontal cortex; EBA = extrastriate body area; IPL = inferior parietal lobe; IPS = intraparietal sulcus; PCUN = precuneus; PLOTC = parietal occipito-temporal cortex; VLPFC = ventrolateral prefrontal cortex.

**Table 2.** Whole Brain Group level statistics of the classification accuracies of explicit vs. implicit conditions. Results produced by the searchlight GNB tested against chance level at q(FDR) < .05 and cluster size corrected (min. cluster size threshold = 176). The values of the peak voxel of each surviving cluster is reported. The degrees of freedom were 19 and p-values were less than .001. The labels in bold represent the clusters resulting from the whole brain statistical map. Regions indicated in normal font are manually defined subregions of the main clusters displayed for completeness.

| **Brain Regions** | **L/R** | **x** | **y** | **z** | **t(19)** |
| --- | --- | --- | --- | --- | --- |
| **Parietal occipitotemporal cortex** |  |  |  |  |  |
| Extrastriate body area | R | 54 | -59 | -5 | 7.207** |
|  | L | -44 | -66 | 1 | 9.531*** |
| Inferior parietal lobule | R | 53 | -49 | 25 | 7.448*** |
|  | L | -53 | -49 | 25 | 4.957* |
| Intraparietal sulcus | R | 35 | -73 | 36 | 8.051*** |
|  | L | -27 | -77 | 36 | 6.918** |
| Precuneus | L | -6 | -68 | 59 | 7.283** |
| **Ventrolateral prefrontal cortex** | R | 48 | 14 | 26 | 10.375*** |
| **Dorsomedial frontal cortex** | L | -12 | 9 | 53 | 6.229** |
| **Cerebellum** | L | -10 | -84 | -30 | 5.769* |

\* p<.0001
\*\* p<.00001
\*\*\* p<.000001

### Interstimulus decoding

The 20 bodies of the stimulus set differed in a number of ways: besides the before mentioned categories of emotion and skin color, there were also person-specific variations in the details of the body pose (e.g. anger could be expressed in a different way between stimuli). This raises the question of whether these fine-grained variations in pose are part of what is encoded in body sensitive cortex. In order to check whether these differences were also reflected in the MVPs, a GNB classifier was trained to classify the 20 affective bodies. As discussed in the univariate analysis (see Materials and Methods) for 7 participants the trial set was incomplete (12 unique stimuli out of 20), therefore they were excluded from this analysis. A group two-tailed t-test against chance level was performed and the resulting t-map showed significant above chance classification accuracy (at q(FDR) <0.05), in cerebellum (t(12) = 6.84, p < .001), bilateral inferior occipital gyrus (IOG) (right t(12) = 5.84, p < .001, left t(12) = 7.12, p < .001), fusiform gyrus (FG) (t(12) = 5.62, p < .001), primary visual cortex (V1) (t(12) = 4.61, p < .0018) (see Fig. 4).

**Figure 4.**
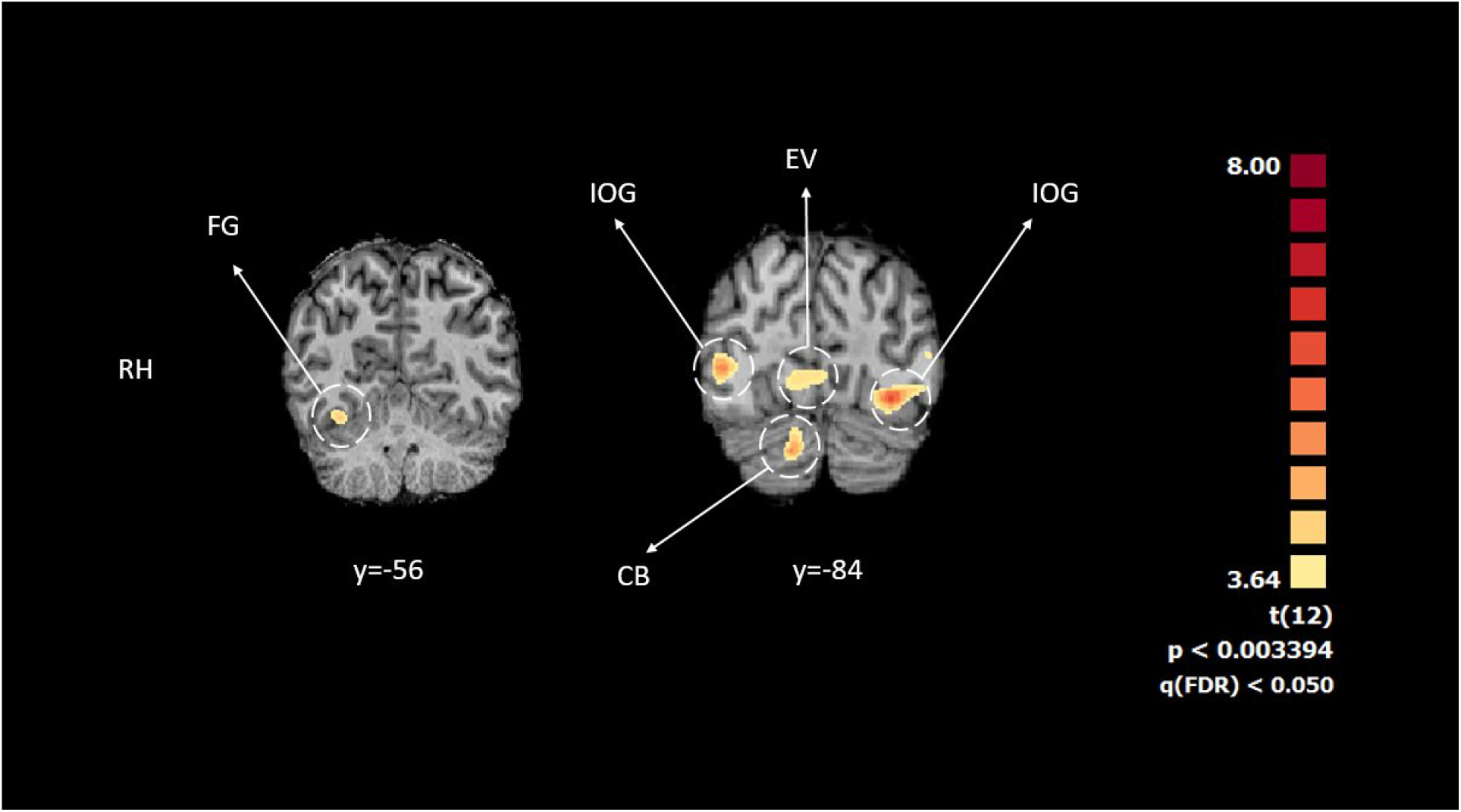
GNB decoding results for all 20 expressive body stimuli. Above chances classification accuracies produced by the searchlight GNB, q(FDR) < .05 for the interstimulus differences are shown. The color map indicates the t-value of the test against chance level accuracy. It is worth noting that IOG is different from EBA here, as the latter is located more anterior in the brain (see Table 2). Abbreviations: CB =cerebellum; EV =early visual cortex; FG =fusiform gyrus; IOG =inferior occipital gyrus.

### Whole brain RSA of intra-versus inter-conditions similarities analysis

In order to determine condition specific (task, emotion, skin) differences in the neural RDMs, we computed for each subject a task specific two sample t-test of intra-condition similarities (e.g. happy-happy, black-black, explicit-explicit) against inter-condition similarities (e.g. angry-happy, black-white, explicit-implicit). When analyzing MVP similarities within the tasks (intra) and between the tasks (inter) we found higher intra-task similarities in bilateral VLPFC, right superior temporal sulcus (STS), bilateral IPS and DMPFC (see Table 3). Here also, we found substantial overlap of results with the GLM and GNB analysis, see Fig. 5b.

**Figure 5.**
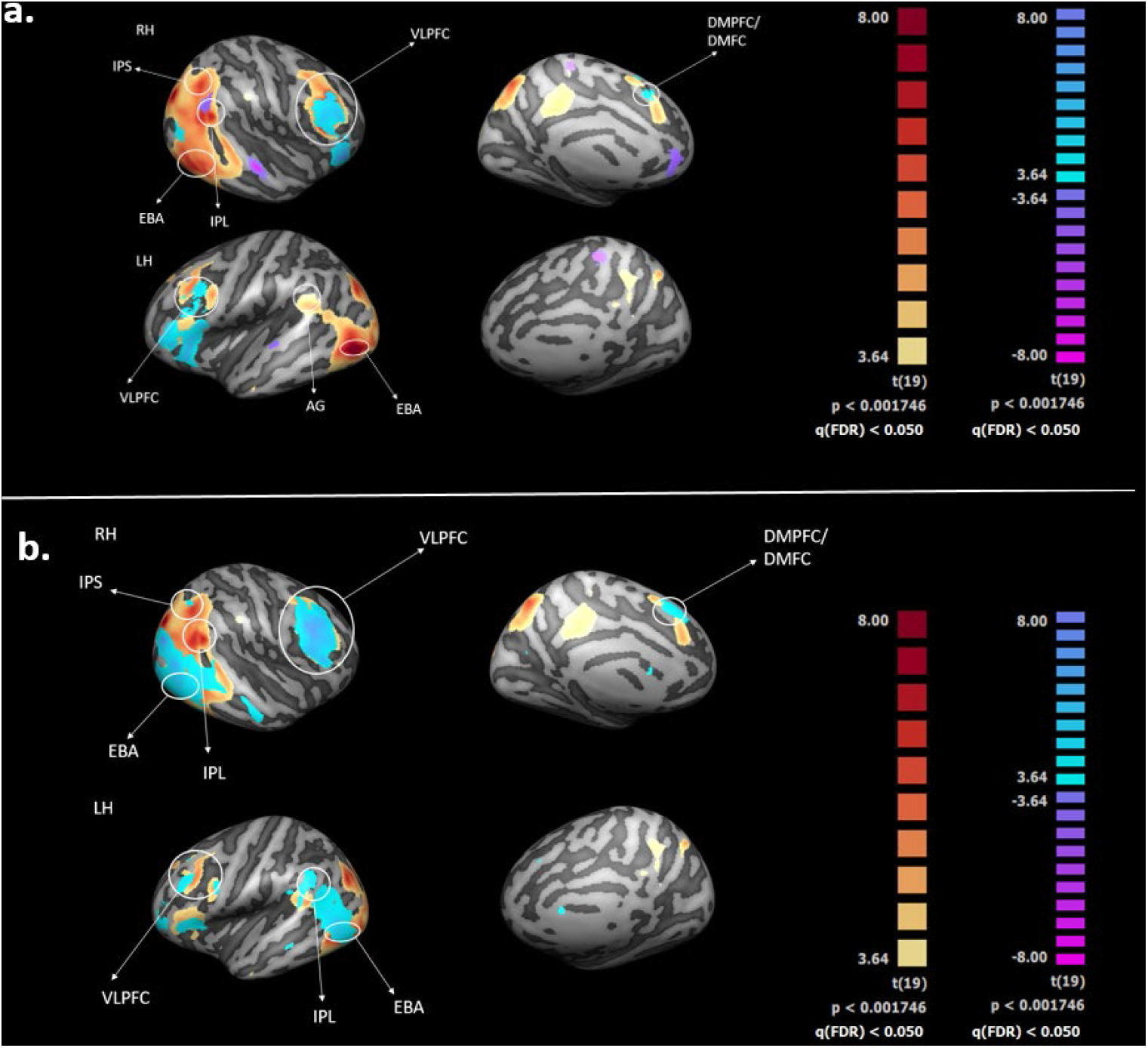
**(a): Whole Brain MVPA and Univariate results overlap:** Combined map of the results of tasks comparison (emotions vs. shapes), mapped to and overlaid on the inflated group average cortical surface, for searchlight GNB (red/yellow) and univariate (blue/purple) results showing the extent of the overlap in RH for VLPFC, IPL and EBA. Abbreviations: AG = angular gyrus, DMFC = dorsomedial frontal cortex; EBA = extrastriate body area; IPL = inferior parietal lobule; VLPFC = ventrolateral prefrontal cortex. **(b): Overlap between GNB results (explicit vs implicit) and intra/inter condition similarities between the explicit and the implicit task.** Shown in light blue/purple are the resulting areas of the inter/intra task similarities analysis (task specific differences in the neural RDMs) at q(FDR) < .05. In order to qualitatively assess the overlap, we superimposed this map on the above chance classification accuracies map produced by the searchlight GNB for the explicit vs implicit expression recognition task (as in panel *a* of this figure), q(FDR) < .05, shown in red/yellow. The positive values (light blue) represent regions which show a higher intra-tasks similarity. Abbreviations: DMFC = dorsomedial frontal cortex; DMPFC = dorsomedial prefrontal cortex; EBA = extrastriate body area; IPL = inferior parietal lobe; IPS = intraparietal sulcus; VLPFC = ventrolateral prefrontal cortex.

**Table 3.**
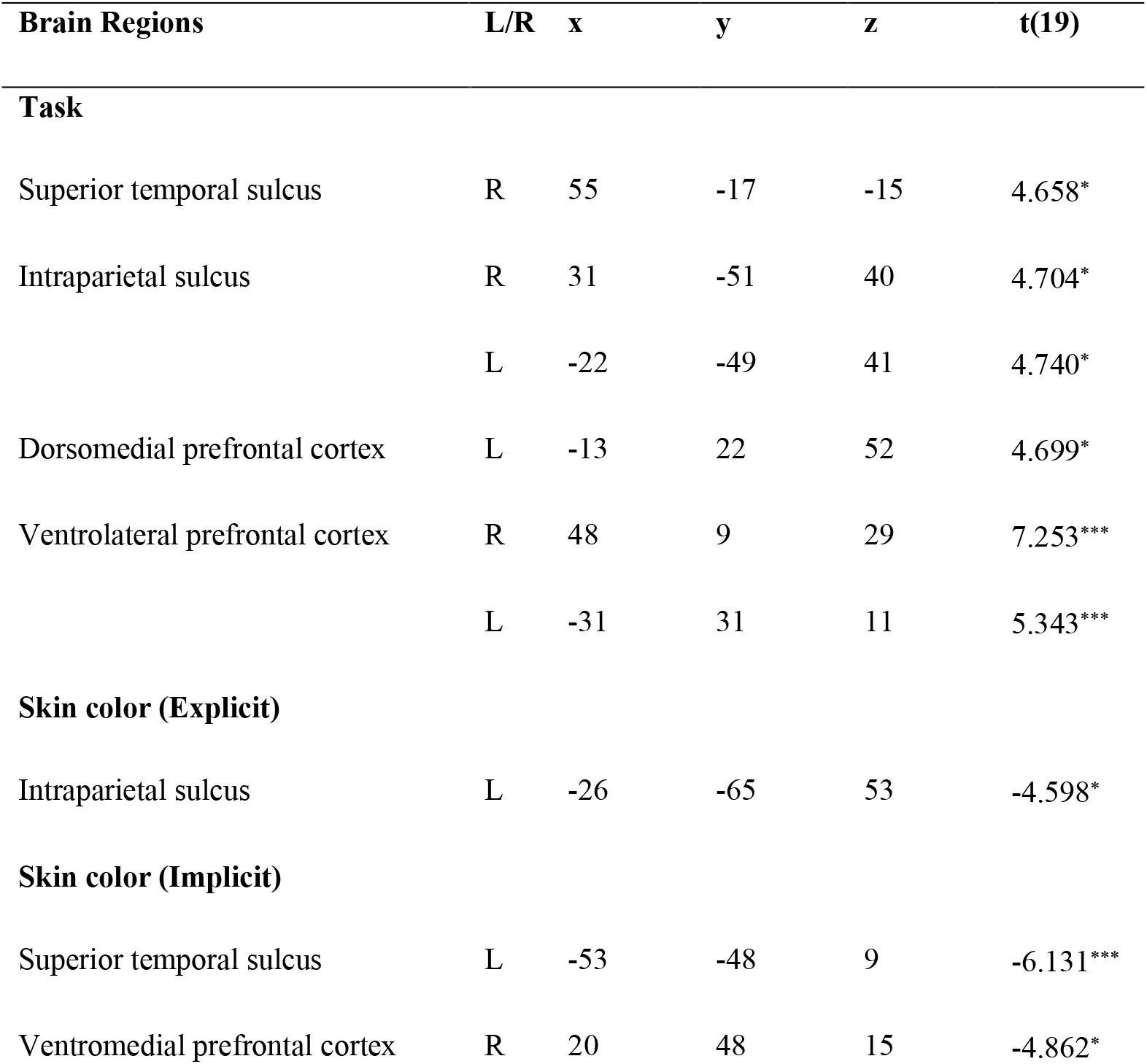
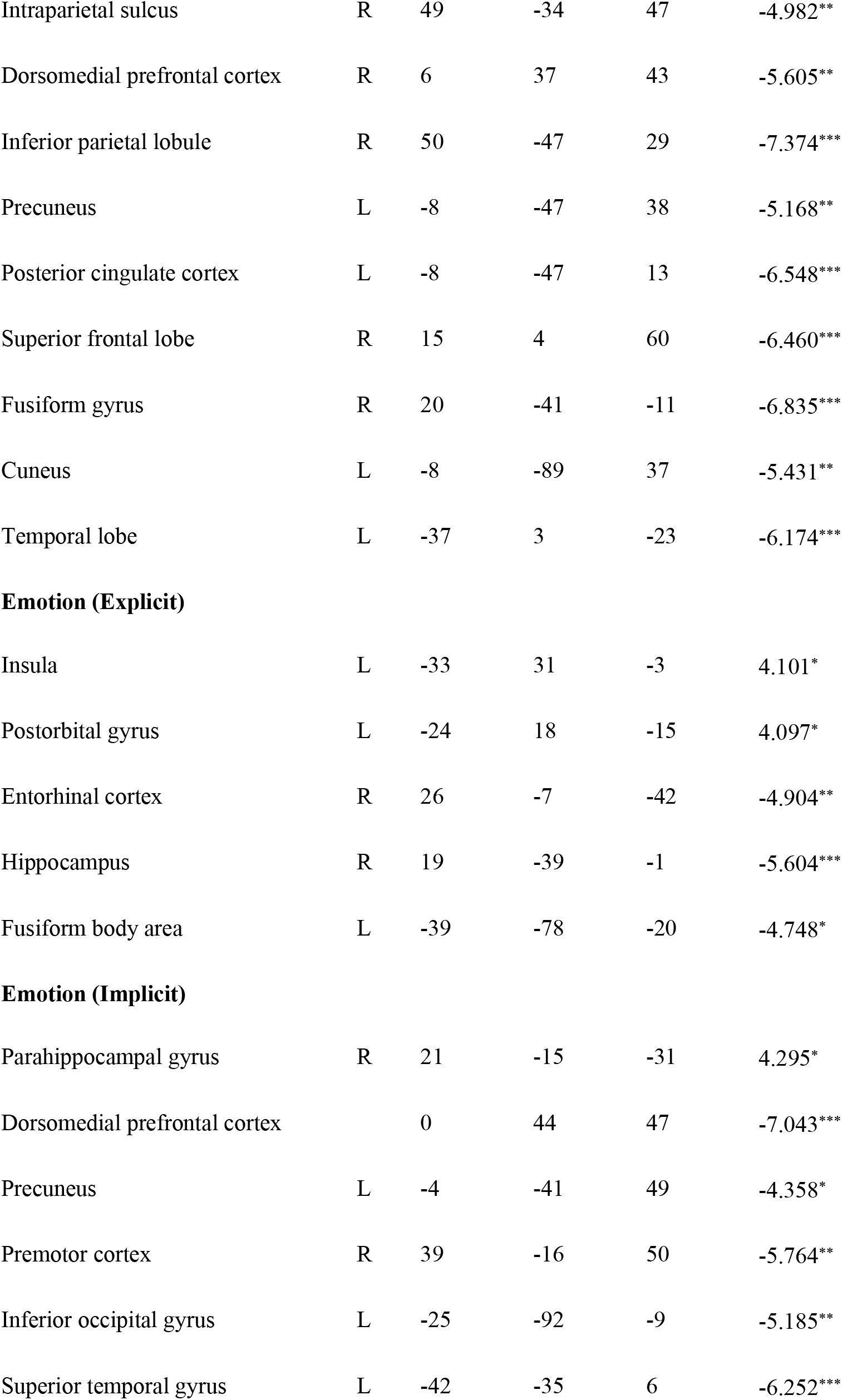
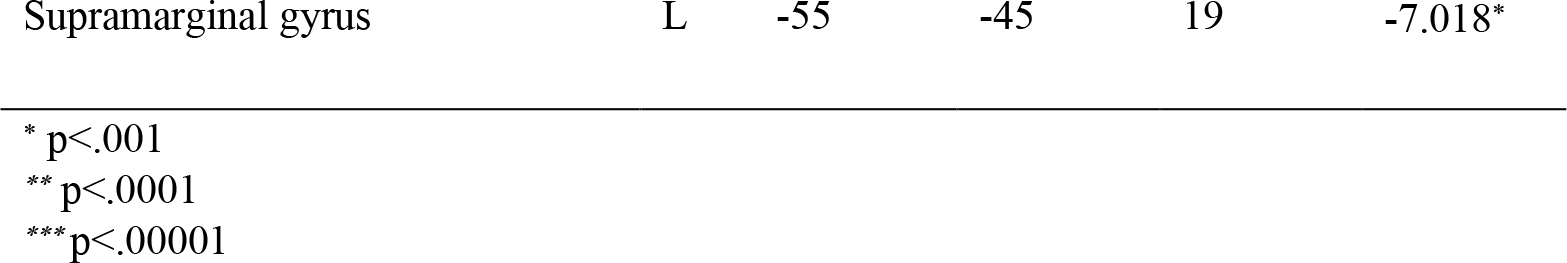
Whole Brain Group level statistics of RSA’s condition specific (task, emotion, skin) effects of multivoxel similarities, at q(FDR) < .05. The table shows the brain regions presenting a higher intra-condition similarity (e.g. happy-happy, black-black, explicit-explicit) (t>0) and those with higher inter-condition similarities (e.g. angry-happy, black-white, explicit-implicit) (t<0). The t values refer to the peak voxel of each surviving cluster. The degrees of freedom were 19 and p-values were less than .001.

We extracted responses to emotion and skin color conditions within the two tasks in order to find regions with higher intra-conditions similarities (i.e. similarity between happy-happy > similarity between happy-angry) and vice versa regions with higher inter-conditions similarity (i.e. similarity between happy-angry > similarity between happy-happy). In the explicit emotion recognition task at q(FDR) = .05, higher similarities between same emotions (higher intra-similarities, happy-happy, angry-angry) were seen in left insula, left post-orbital gyrus, whereas higher similarities between different emotions (higher inter-similarities, happy - angry) were found in right entorhinal cortex, right hippocampus, left FBA (see Fig. 6 and Table 3).

**Figure 6.**
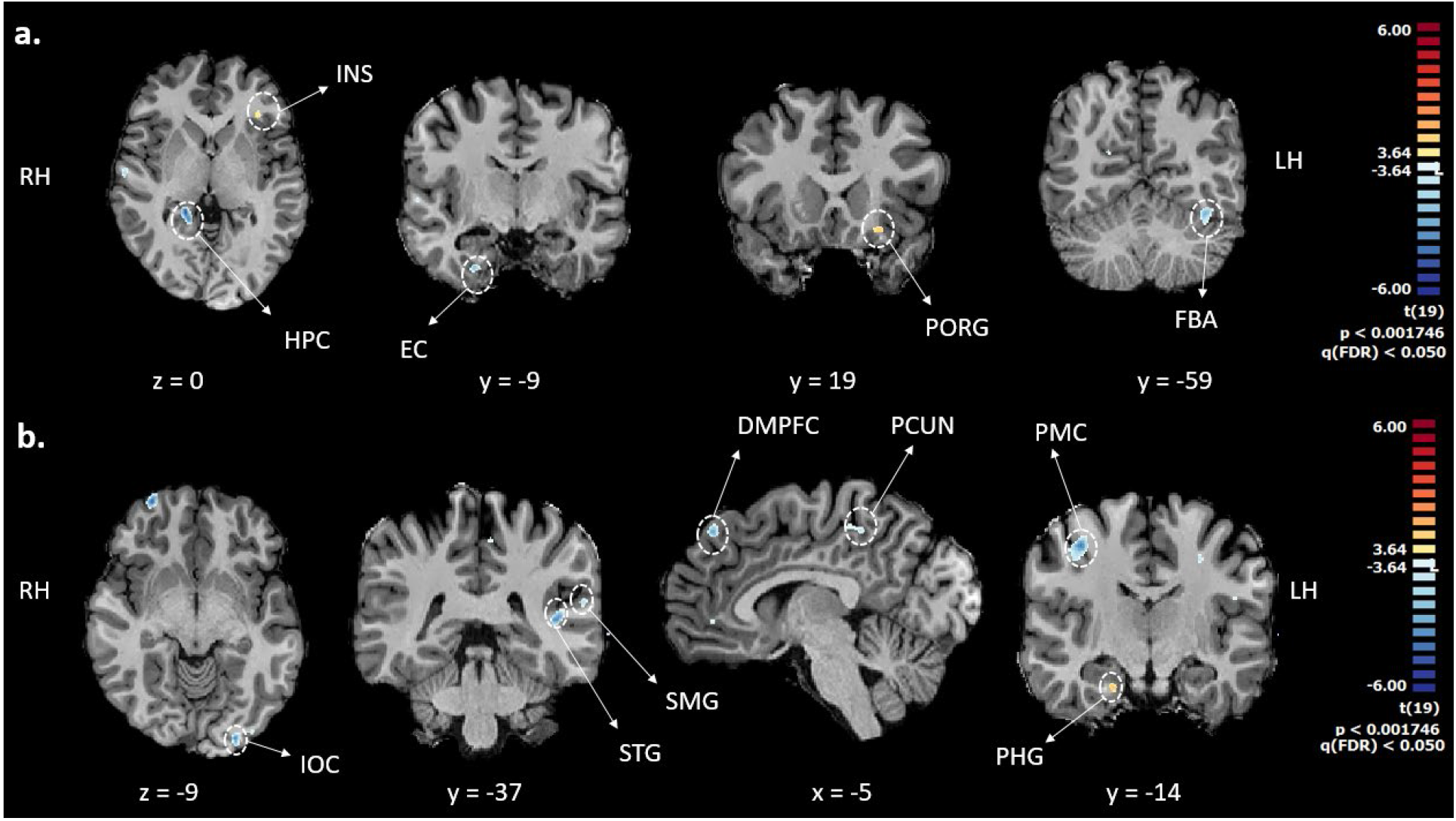
Inter/Intra emotion similarities analysis: Task specific results for affective body postures (angry, happy) in explicit (a) and implicit (b) emotion recognition. Group results of the two-sample t-test between intra-emotions similarities against inter-emotions similarities at q(FDR) < .05. Panel a (explicit task) and panel b (implicit task) represent brain regions in which neural RDMs for same emotions are more similar than the neural patterns for different emotions (red) and vice versa (blue). Abbreviations: EC = entorhinal cortex; HPC = hippocampus; INS = insula; DMPFC = medial prefrontal cortex; PMC = premotor cortex; PORG = post-orbital gyrus.

In the implicit emotion recognition task, higher similarities were found between same emotions (higher intra-similarities) in right parahippocampal gyrus, whereas higher similarities between different emotions (higher inter-similarities) were found in dorsomedial prefrontal cortex, left precuneus, right premotor cortex, left inferior occipital gyrus, left superior temporal gyrus, left supramarginal gyrus (see Fig. 6 and Table 3).

For the explicit task, higher similarities between different skin colors (higher inter-similarities, black-black, white-white) were found in left IPS. Similarly, in the implicit task higher similarities between different skin colors (higher inter-similarities, black-white) were found for DMPFC, ventromedial prefrontal cortex (VMPFC), left precuneus, right IPS, right IPL, right superior frontal lobe (SFL), left temporal lobe, left cuneus, left PCC, right FG, left PSTS (see Fig. 7 and Table 3).

**Figure 7.**
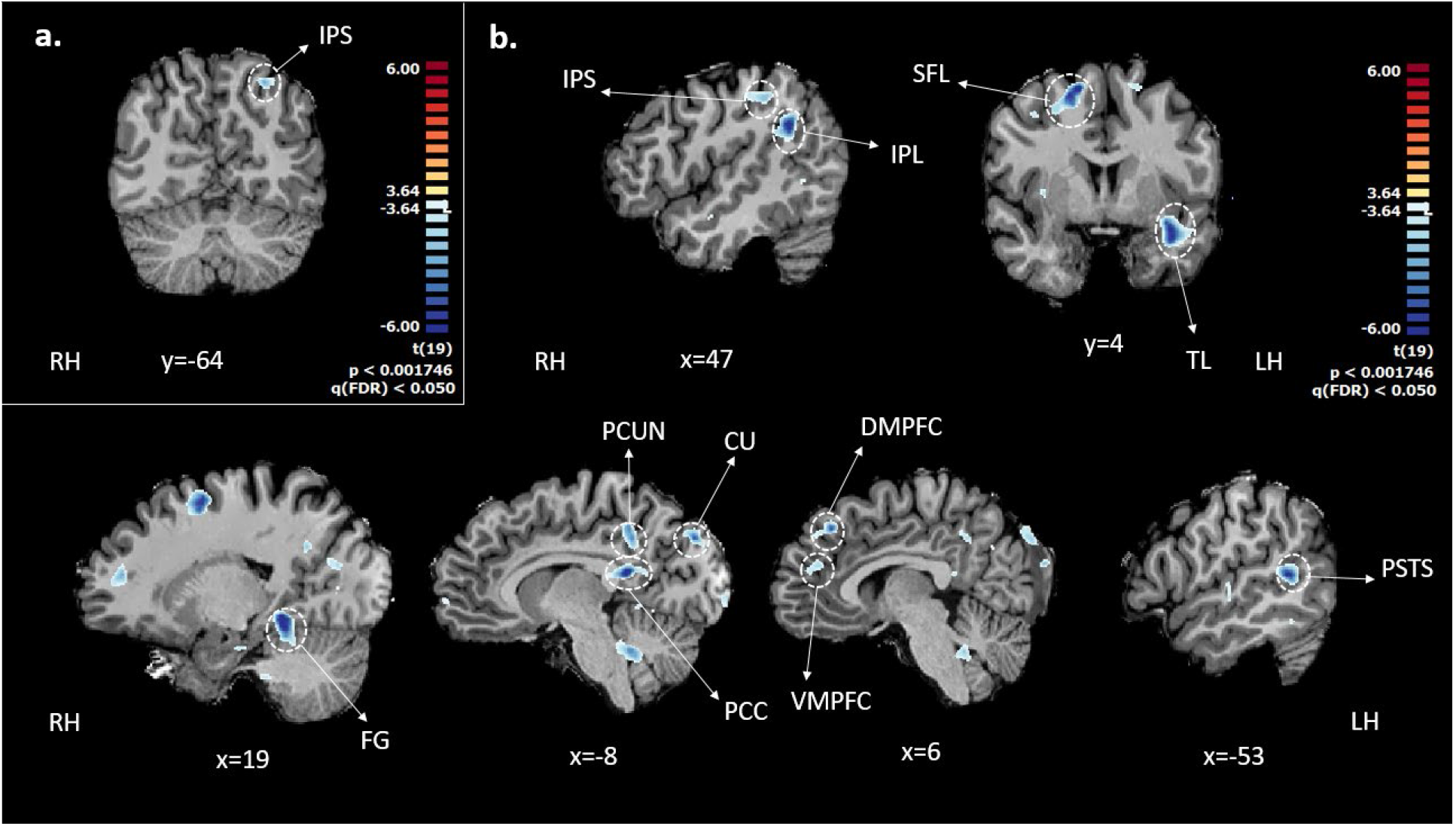
Inter/Intra condition similarities analysis: Task specific results for skin colors (black, white) in explicit (a) and implicit (b) emotion recognition. Group results of the two-sample t-test between intra-condition (e.g. black-black) similarities against inter-conditions similarities (e.g. black-white) at q(FDR) < .05. Panel (a) and panel (b) represent brain region in which neural RDMs for same emotions are more similar than the neural patterns for different emotions (red) and vice versa (blue) for the explicit task and implicit task respectively. Abbreviations: CU = cuneus; DMPFC = dorsomedial prefrontal cortex; FG = fusiform gyrus; IPL = inferior parietal lobule; IPS = intraparietal sulcus; VMPFC = medial prefrontal cortex; PCC = posterior cingulate cortex; PCUN = precuneus; PSTS = posterior superior temporal gyrus; SFL = superior frontal lobe; TL = temporal lobe.

### Region of Interest Analysis

The analyses on task effect (univariate GLM, multivariate GNB) revealed convergent results spanning a number of anatomical regions (Fig. 3), e.g. VLPFC, IPL and LOTC (including EBA). To gain a more detailed insight into the responses in these regions, we defined ROIs via a 5-fold cross-validation procedure (see Material and Methods). The ROIs differed in size and location (see Table S8) across folds, however as shown in Fig. 8 the extent of the overlap was consistent across folds. For the contrast considered (explicit vs. implicit task decoding) we extracted within each fold the peak voxel of each ROI from the training set by setting a statistical threshold q(FDR) < .01. This revealed bilateral EBA, right IPL, right VLPFC, precuneus, and right IPS, see Table S8.

**Figure 8.**
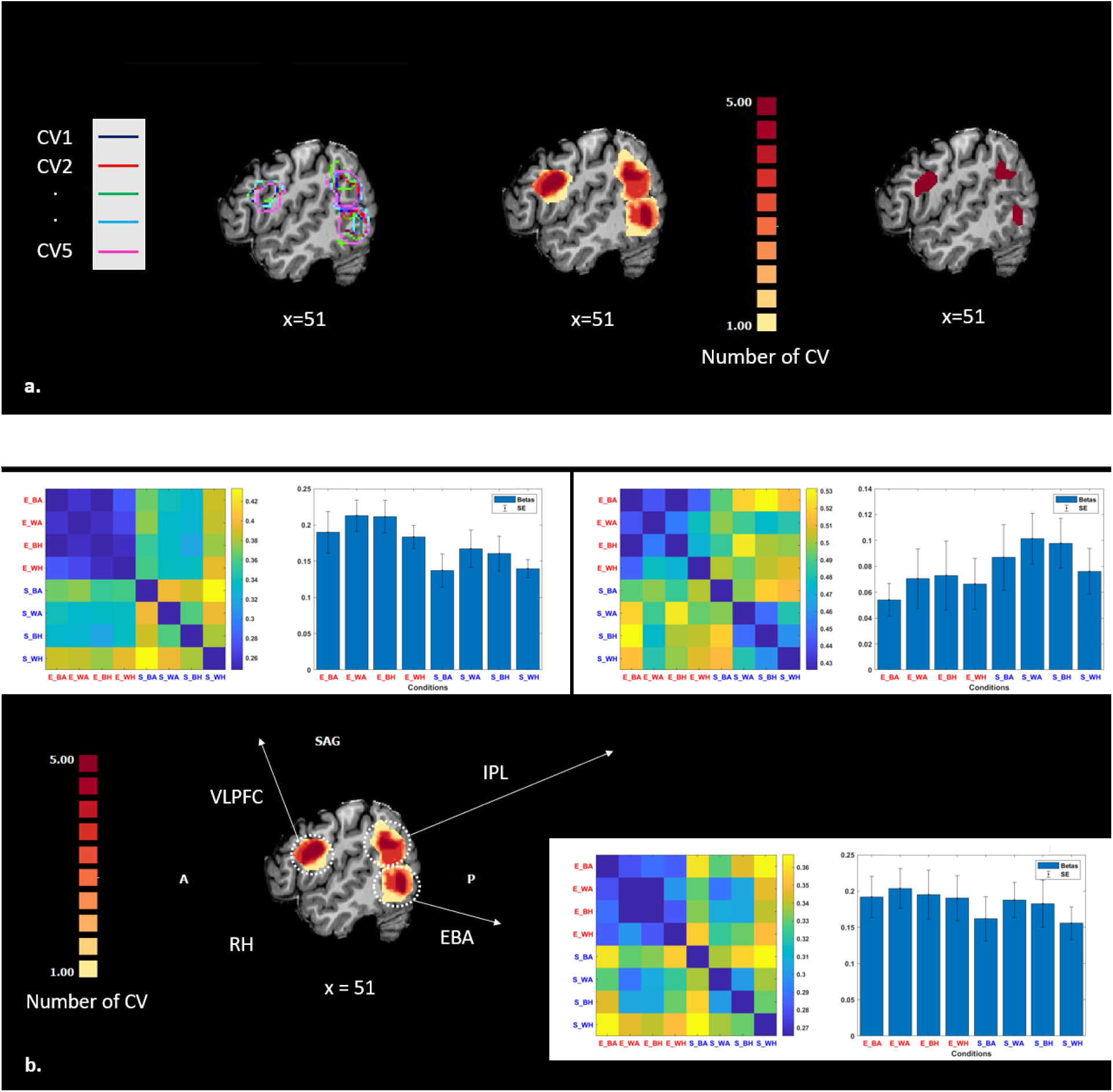
**(a): ROIs identification from GNB task decoding (explicit vs. implicit) accuracies maps and overlap across folds.** In the left panel we show the contour of the regions identified during the 5-fold cross-validation for the ROIs under examination: right EBA, right VLPFC and right IPL, each color identifies a specific fold. In the middle panel we show a gradient map of the overlapping voxels across folds from yellow (no overlap across folds) to dark red (full overlap: voxel selected in all 5 folds). In the right panel we plot the regions where we found complete overlap across folds (same voxels shown in the middle map in dark red). **(b): Details of the responses from the ROIs identified during the cross-validation procedure, RDM and beta plots at the category level of each ROIs are shown.** The different ROIs were defined using a 5-fold cross-validation on the task based decoding (explicit vs. implicit) accuracies maps computed at a single subject level (see Material and Methods). We display the RDMs and beta plot (averaged across folds) on the clusters which show the overlap between folds from yellow (no overlap across folds or voxel selected only in 1 fold) to dark red (full overlap across folds or voxel selected in all the folds), as shown in panel (a) (middle). For each fold and for each resulting ROI (panel (a), left) we computed the RDMs and beta values by extracting the activity pattern of each subject which was left-out during the procedure of ROI definition. Within each fold RDMs and beta values were averaged across participants. Ultimately, the beta and RDMs plots displayed in panel (b) where defined by averaging across folds the RDMs and the beta values computed within each instance of the cross-validation. In the beta panel we plot the mean plus standard error averaged across folds of the 8 conditions. Abbreviations: EBA = extrastriate body area; IPL = inferior parietal lobe; VLPFC = ventrolateral prefrontal cortex.

As shown in Fig. 8b, the neural RDMs of the EBA and VLPFC ROIs show a similar structure, in particular in the explicit task conditions (upper left half of the RDM), whereas this effect is absent in the implicit conditions (bottom right half of the RDM). While the MVPs of the other regions (see supplementary material, Figs S1 and S2) produce RDMs which present effects (similarities or dissimilarities) within conditions or activation levels, they do not show the clear pattern found for right EBA and VLPFC. In order to check for activation differences between the two tasks, we performed a t-test between beta values averaged across tasks within each cross-validation, this revealed higher activation for the explicit task in VLPFC (t(4) = 4.69, p=.009) and higher activation for the implicit task in IPL (t(4) = −2.74, p = .051).

## Discussion

In the present study we measured the representational dynamics of explicit and implicit body expression perception and identified the brain areas that are critical for the distinction between the two tasks. Our results revealed three main findings. First, the difference between explicit and the implicit body expression processing can be decoded with high accuracy in right EBA, VLFPC and IPL. Second, the brain activity associated with explicit recognition in these areas is not emotion specific. Third, in contrast, some specific effects for different emotions are observed in the implicit condition. In the sections below we discuss these findings and propose that taken together these findings suggest that the way in which object category, stimulus attributes and action are represented is dynamically organized by the requirements of each task and contributes to clarifying the functional role of body areas.

### Similar task specificity across high-level visual cortex, EBA, VLPFC and IPL

The first major result of our study is that there are three areas where the difference between naming the expression or naming a shape while ignoring the expression can be decoded with high accuracy as seen in highly similar responses for all conditions in the explicit task. Our results are consistent with previous studies that have reported task specific activity in VLPFC (Bracci, Daniels, and Op de Beeck 2017; Bugatus, Weiner, and Grill-Spector 2017; Xu and Vaziri-Pashkam 2019; Haxby, Connolly, and Guntupalli 2014; Kriegeskorte et al. 2008; Pichon, de Gelder, and Grezes 2009) and is consistent with role of cognitive and affective control attributed to VLPFC (Szczepanski and Knight 2014). Task sensitive activity level in higher visual areas is more debated and was found in some but not in other earlier studies. A previous study (Bugatus, Weiner, and Grill-Spector 2017) found that during either a working memory, oddball or selective attention task, the task effect was limited to VLPFC and not seen in high-level visual cortex where responses were more driven by stimulus category than by the task demands, in line with classical view on category specific areas. One explanation for the same task effect seen in EBA and VLPFC here is that VLPFC contains flexible category representations (here body selective neurons) that are mobilized when the task requires it (Bugatus, Weiner, and Grill-Spector 2017). However, while this may explain the observed task sensitivity to body expression categorization in VLPFC, it does not address the associated task sensitivity in right EBA. An alternative explanation that would clarify that similar task effects are found in EBA and VLPFC is that the explicit task effect we see here reflects selective attention. Body category perception driven by selective attention to the expression might then have a region-general effect across EBA and VLPFC. This is in agreement with studies showing that selective attention alters distributed category representations across cortex, and particularly in high-level visual cortex and also in VLPFC (Cukur et al. 2013; Peelen, Fei-Fei, and Kastner 2009; Shahdloo, Çelik, and Çukur 2020). These studies found effects of selective attention-based increases in category representation areas for the preferred category in visual search tasks.

Our results are consistent with this to some extent as selective attention to the body expressions in the explicit task may boost body category representation in EBA consistent with findings that emotional expression increases EBA activity (de Gelder and Poyo Solanas 2021; Peelen et al. 2007). But then such an attention-based activity increase should possibly be visible in FBA as well. On the other hand, there is evidence that category selective mechanisms in visual object areas operate outside selective attention, a process attributed to neural mechanism for attentional selection enshrined in the category selective area (Peelen, Fei-Fei, and Kastner 2009). This would lead one to expect little difference between activity in body areas between the explicit and the implicit task, contrary to what is found here. Unless, indeed as also suggested by the literature, EBA plays a more important role in expression perception than FBA.

### Task dynamics, body and body representation in EBA

EBA and FBA are commonly viewed as ventral stream areas associated with body representation but their respective functions are not yet clear nor is their anatomy well understood (Weiner and Grill-Spector 2012). Whole body perception is attributed more to FBA than to the EBA which is seen as more involved in body parts (Downing et al. 2001; Peelen and Downing 2007). Few studies have yet investigated the specific functional roles of FBA and EBA either in expression perception or in relation to task demands and available studies find no clear differences in their functional role for expression and task sensitivity. Our results contribute to clarifying this situation.

Considering more specific functions of category sensitivity, a current view is that EBA encodes details pertaining to the shape, posture and position of the body and does not directly contribute to high level percepts of identity, emotion or action that are potential functions of FBA through its connections with other areas (Downing and Peelen 2011). However, studies on body expressions have most often reported involvement of both EBA and FBA with the activity pattern varying with the specific expression considered but without any clear understanding of the respective functions (Costantini et al. 2005; Marsh et al. 2010; Moro et al. 2008; Pichon, de Gelder, and Grezes 2012; Saxe, Jamal, and Powell 2006; de Gelder, de Borst, and Watson 2015; Tamietto et al. 2015; Van den Stock et al. 2015).

Recent evidence projects a more detailed view on EBA and how it could contribute differentially to body and body expression perception which is consistent with our present findings. First, an investigation aimed at sorting out the function of EBA and adjacent MT+ reported a double dissociation. TMS over EBA disrupting performance in the form discrimination task significantly more than TMS over pSTS, and *vice-versa* for the motion discrimination task (Vangeneugden et al. 2014). Additionally, (Zimmermann et al. 2016) showed that early disrupting of neuronal processing in EBA during action planning, causes alterations in goal-oriented motor behavior. Second, in support of the differences found here, EBA and FBA show a very different profile of anatomical connectivity with other brain areas, notably with parietal areas (Zimmermann et al. 2018). Third, EBA is a complex area with important subdivisions (Weiner and Grill-Spector 2011) possibly coding different features of whole body images. In line with this, (Ross 2014) propose to dissociate the EBA-MT+ area as this would profile EBA more clearly as the area coding body form and clarify functional differences between EBA and FBA. In a recent study investigating detailed features of body expressions and how they are represented in the brain, major differences were found in the functional role of EBA and FBA when studied at the feature coding level (Poyo Solanas, Vaessen, and de Gelder 2020b). EBA and FBA also showed tuning to postural features of different expressions. However, the feature representation in EBA was very dissimilar to that of FBA. Similar feature representation to that seen in EBA was found in SMG, pSTS, pIPS and the inferior frontal cortex but not in FBA (Poyo Solanas, Vaessen, and de Gelder 2020b). When such findings targeting function descriptions at the feature level accumulate, more detailed hypotheses about task effects become feasible.

Another possibility is that the effects observed in EBA reflect recognition of the body expression perception of only a body part like the hands and not on the whole body. Recent evidence shows that the hands are more informative for certain emotions, including anger images used here (Poyo Solanas, Vaessen, and de Gelder 2020a; Kret and de Gelder 2012; Kret et al. 2017; Ross and Flack 2020). Concerning representation in the brain (Taylor, Wiggett, and Downing 2007) found that bilateral EBA showed a preference for individual body parts such as the hands and fingers while FBA showed a preference for the whole body. (Bracci et al. 2010) showed selective response to hands over other body parts in left EBA. Our results in the explicit condition revealed right EBA instead. Since our study used whole body stimuli and not body parts, we cannot directly address this possibility. But the position of the fixation was intended to counter part based recognition. Furthermore, we did not find emotion specific activity in EBA in the explicit condition as might have been expected if explicit recognition responses would be based on noticing hand position which is more indicative for. Furthermore, as can be seen from the sample images (Fig. 1b), there is some variability in hand position within the same category while the overall configuration is similar. Nevertheless, overall configuration is known to play a crucial role in body like in face perception (Stekelenburg and de Gelder 2004).

### Task decoding and the role of IPL

Besides EBA and in VLPFC, we are also able to decode the difference between the tasks in IPL, albeit less clearly and importantly, with the opposite pattern of higher beta values for the implicit condition. This was also found in the univariate results where IPL is more active in the implicit task. IPL is a hub structure and is involved in at least four networks (the frontoparietal, default mode, cingulo-opercular and ventral attention network (Igelström and Graziano 2017). Previous studies provided clear evidence for the role played by IPL in body and emotional perception. Emotion-specific activation within parietal cortex was found for face stimuli (Grezes, Pichon, and de Gelder 2007; Kitada et al. 2010; Sarkheil et al. 2013) and for body stimuli (de Gelder et al. 2004; Goldberg et al. 2015; Goldberg, Preminger, and Malach 2014; Kana and Travers 2012). Significant activity was elicited in IPL for the contrast bodies expressing fear or happiness (Poyo Solanas et al. 2018). We argued previously that IPL may play the role of a hub where emotion perception is transitioned into an action response (Engelen et al. 2018). IPL receives input from the visual system (Caspers et al. 2011) and has connections to pre-motor cortex involved in action preparation (Hoshi and Tanji 2007; Makris et al. 2005; Mars et al. 2011).

Higher activity in IPL in the implicit task fits the role of IPL in action representation and its involvement in the transition to action preparation (Engelen et al. 2018). Explicit emotion recognition is a cognitive task and in the course of using verbal labels action tendencies triggered by the stimuli tend to be suppressed, which may be reflected in lower IPL activity (Engelen et al. 2015; Igelström and Graziano 2017). Consistent with this and as argued above, there is no difference between the emotion conditions in the explicit task while there is a suggestion of this in the implicit task (but this is not significant).

### The role of VLPFC

Similar to the results for right EBA we found that activity in right VLPFC allows decoding the task difference, again with significantly higher beta values for the explicit task and with no difference between the expression conditions. In the whole-brain RSA, VLPFC showed higher intra-task similarity (higher similarity for same task) (see Fig. 5 and Table 3), consistent with the pattern of similarities we found in the RDMs during the ROIs analysis (see Fig. 8). The literature suggests different explanations for the role of VLPFC. One is its role in attention and decision making, another one the possibility that VLPFC contains object category representations and finally, a role of VLPFC in regulating affective processes. The latter alternative is best supported by the pattern of results.

A familiar function of VLPFC is related to theories of PFC as predominantly involved in attention and decision processes (Duncan 2001, 2010) and it associates VLPFC activity with increased task demands (Crittenden and Duncan 2014). But our two tasks were designed to be very similar in difficulty and in cognitive demands and required a simple forced choice between two alternative responses. Under these circumstances one would not expect a task related difference in VLPFC and indeed accuracies are near 100%. Similarly, attention is known to be triggered selectively by some body emotion expressions (eg. fear) more than others (de Gelder, Hortensius, and Tamietto 2012; Tamietto et al. 2015; Zhan, Goebel, and de Gelder 2018). Yet we do not observe a difference between the emotions as would be expected it the VLPFC activity corresponded to endogenous attention. This speaks against the notion that VLPFC activity here reflects an effect of attention. A second explanation is that VLPFC activity reflects a task effect and not an attention effect (Bugatus, Weiner, and Grill-Spector 2017) based on the notion that VLPFC is the final stage of high level vision in the ventral pathway involved in categorization (McKee et al. 2014; Bugatus, Weiner, and Grill-Spector 2017; Cukur et al. 2013; Peelen, Fei-Fei, and Kastner 2009). However, those studies used a number of different object categories unlike the present study only using bodies and where the explicit task was expression recognition. This makes it unlikely that the present role of VLPFC reflects a task effect based on category selectivity.

In contrast with those two alternatives our results best support the notion that VLPFC is involved in suppression of emotion related processes that are automatically triggered by presentation of emotional stimuli. Previous studies have shown that the VLPFC is involved in downregulating emotion responses presumably based on its structural and functional connectivity to the amygdala (Wager 2008). TMS directed on VLPFC, interrupted processing of emotional facial expressions (Chick et al. 2019). The fact that beta values are higher in VLPFC for explicit recognition conditions is consistent with this explanation.

### Explicit vs implicit representation of emotions

A first finding of the RSA is that decoding accuracies for emotion were overall low and did not differ between the emotion and the shape task. In the Intra/Inter RDMs similarities analysis (Fig. 6,7) specifically looking for emotion condition effects, we did observe an overall pattern of task and emotion representation dynamics. Overall, we find similarities and differences between the emotion conditions for the two tasks. For the explicit emotion recognition task, higher similarities between same emotions were seen in left insula and left post-orbital gyrus. Interestingly, these areas are found when body expressions are viewed consciously but not when they are unattended or neglected (Tamietto et al. 2015; Salomon et al. 2016). For the implicit emotion recognition task, higher intra emotion similarities were found in right parahippocampal gyrus, which may reflect that processing expressions involves memory similarly for both expressions. For the explicit task, higher similarities between different emotions presumably representing what is common to different emotions, were found in right entorhinal cortex, right hippocampus and left FBA. Concerning the latter, this suggest that FBA is involved in expression recognition but does not contribute to specific expression coding. In contrast, in the implicit task higher similarities between different emotions were found in medial prefrontal cortex, left precuneus, left premotor cortex, right inferior occipital gyrus, right superior temporal gyrus and right supramarginal gyrus. Interestingly, the latter are all areas known from studies that used passive viewing or oddball tasks and not emotion labeling or explicit recognition (de Gelder et al. 2004; Grezes, Pichon, and de Gelder 2007; Goldberg et al. 2015).

However, we can relate the EBA and VLPFC results to the role of IPL in action perception and preparation as discussed above. The finding of task sensitive activity in IPL suggests that the higher similarities in the explicit emotion task for VLPFC and EBA are not just independently reflecting stimulus/task settings and higher activation level in the explicit emotion task. The combination of higher activation in EBA and VLPFC and lower activation in IPL suggests connections between these three areas with VLPFC possibly influencing EBA positively and IPL negatively (Ongur and Price 2000; Goldman-Rakic 1996; Ong, Stohler, and Herr 2019; Craig 2009; Tamietto et al. 2015). For explicit recognition of the body expression, category representation would be strengthened while emotion action related information would be suppressed. Further studies using connectivity measures are needed to support this hypothesis.

It is also worth noting that the amygdalae were not among the areas we found to be important for task decoding. The GNB classifier used for the analysis was trained to find regions with large differences in the MVPs for the explicit and the implicit task and did not reveal the amygdalae. Many studies have argued that the amygdala is activated for facial and body expressions of fear, anger of happy expressions and that activity can be lower under implicit viewing conditions (de Gelder, Hortensius, and Tamietto 2012; di Pellegrino, Rafal, and Tipper 2005; Habel et al. 2007; Lieberman et al. 2007). The fact that this difference does not emerge here for the amygdalae may have different reasons. First, the literature is not clear on this issue as a reduced amygdalae involvement is not systematically reported. Second, this result may obviously be related to poor SNR in that area. Third, it is difficult to generalize effects at the level of the whole amygdalae, given their multiple nuclei with very different functions. On the other hand, we do find task and expression differences in areas that are known to be functionally connected to the amygdalae, most importantly the IPL. Patients with amygdala damage show decreased connectivity between basolateral amygdalae and prefrontal and temporal areas under conditions of task irrelevant body expression perception but increased connectivity between the same amygdala nucleus and IPL (Hortensius et al. 2017). This might be an indirect signature of a role for amygdalae involvement in the sense that in the implicit task here IPL activity is higher than in the explicit task.

### Limitations and future perspectives

As the present study used two body expressions further research is needed to conclude whether the same pattern of differences between implicit and explicit perception would be observed with different emotional expressions like for example fear or sadness. On the other hand, generalization to other emotions should not be taken to mean that the same pattern is expected across different emotions. It is known from previous studies that stimuli of different emotion expressions behave differently in experiments measuring non-conscious processing like for example when CFS is used (Zhan, Goebel, and de Gelder 2018; Zhan and de Gelder 2019). Importantly, these differences are not expected as long as emotions are viewed as abstract concepts, and emotion perception is a matter of applying abstract concepts (see above), but they are very likely in a naturalistic and behavioral perspective. For example, fear and anger automatically prompt behavioral reactions that sadness does not. Our goal was not to discover a pattern that would generalize across different emotions. This might be expected since we generally observe high recognition accuracy for all basic emotions (de Gelder and Van den Stock 2011), suggesting that similar task related differences would also be found for other emotions. This expectation reflects the traditional concept-based view on emotion perception. High accuracy recognition rates for body expressions do not directly provide evidence for similarity in associated adaptive behavior and underlaying neural processes associated. There are very different views in the literature about the relation between emotion words used in reports of subjective recognition and neurobiological bases of the underlaying processes (Mobbs et al. 2019). Indeed, a widely held view is that the brain decodes emotion stimuli by using higher-order conceptual emotion representations typically used in descriptions of mental states.

Another possible limitation concerns the number of identities. However, the postures display standard expressions that are effortless recognized as can be seen in the behavioral results. And because facial identity information is blurred, individual personal identity of each stimulus is unlikely to impact the results. Given how our stimuli were created, some variability between the postures is to be expected. Actors were instructed to react to a given situation, familiar from daily life. They were not asked to express an emotion and were not given abstract emotion labels. Of course, in daily life the situations they were asked to react to are typically associated with typical emotion labels. Some actors are more expressive than others and this presumably reflects personal style, personality, extroversion. Still variability is limited in the sense that across actors the same body parts are involved. For example, as can be seen from the images in Fig. 1b, anger involves the hands besides also the leg position and the overall posture. So, there is variability in the stimulus set, as there is variability in people’s expressions in daily life. We believe that it is important to note that interindividual variability cannot be well judged with the naked eye and its contribution to the result cannot be assessed reliably by looking at the images. We would need computational models allowing quantitative description and computational analysis of the posture features in order to have an objective assessment of whether variations in feature positions (angle of the arm, direction of the hand etc) matter for how the brain encodes the body postures (de Gelder and Poyo Solanas 2021). An example of such a computational analysis of body features was undertaken for still images (Zhan, Goebel, and de Gelder 2021) and for video images (Poyo Solanas, Vaessen, and de Gelder 2020b).

Another limitation of our study is that the design used does not allow to measure functional relations between the critical areas observed. Further studies using connectivity measures are needed to support our suggested explanation. Finally, is worth noting that while two decades of neuroimaging on the brain correlates of human emotion have not yielded a clear picture of how emotions are represented in the brain (Wager et al. 2015). But relatively few studies have contrasted explicit recognition and implicit perception and the few studies who did so find substantial differences for body expressions (Zhan, Goebel, and de Gelder 2018). Besides the theoretical importance of the distinction, this task contrast is particularly relevant for understanding emotion perception in clinical populations like schizophrenia (Trémeau et al. 2015) and autism (Jones, Lambrechts, and Gaigg 2017; Luckhardt et al. 2017). For example, in studies of autism and schizophrenia it has been reported that implicit measures are more diagnostic than explicit ones (Hajdúk et al. 2019; Luckhardt et al. 2017; Van den Stock et al. 2011). A better understanding of implicit emotion processing as seen in real life routines and explicit recognition as seen in questionnaires may shed new light on clinical findings and provide a rich analytical framework for investigating social cognitive disorders.

## Conclusion

The main purpose of this study was to investigate how explicit and implicit emotion perception tasks affected activity in body category and emotion coding areas and to assess whether the activity patterns would also reflect differences between emotional expression and skin colors. Overall, this result indicates that the similarities found in explicit tasks do not map onto the pattern of the implicit ones and stress the importance of the specific task both when investigating category selectivity and brain correlates of affective processes. The clear task effects seen here also indicate that understanding category and emotion attribute representations may profit from being viewed in the larger context of connectivity between ventral category areas and other areas in the brain.

## Acknowledgements

This work was supported by the European Research Council (ERC) FP7-IDEAS-ERC (Grant agreement number 295673 Emobodies), by the Future and Emerging Technologies (FET) Proactive Programme H2020-EU.1.2.2 (Grant agreement 824160; EnTimeMent) and by the Industrial Leadership Programme H2020-EU.1.2.2 (Grant agreement 825079; MindSpaces). We are grateful to Dr. Federico De Martino, Dr. Agustin Lage Castellanos, Dr. Minye Zhan and Marta Poyo Solanas for valuable comments and suggestions on an earlier version and to anonymous reviewers for very constructive criticism.

