## Supplementary Material for "The representation of bodies in high level visual, prefrontal and inferior parietal cortex varies with explicit vs. implicit expression perception"

**Table S1. Whole Brain Group level univariate results of interaction between task and race**. The table shows the regions where the 3-way ANOVA revealed an interaction between task and skin color factors. The F-map was thresholded at q(FDR) < .05 and cluster size corrected. Peak voxel coordinates (MNI) and corresponding F value of each surviving cluster are reported. The degrees of freedom for the ANOVA 1 and 19. All results were significant at p < .001.

| **Brain Regions** | **L/R** | **x** | **y** | **z** | **F(1,19)** |
|---|---|---|---|---|---|
| Fusiform gyrus | R | 44 | -53 | -15 | 40.037** |
| | L | -46 | -59 | -12 | 43.830** |
| Inferior/Middle occipital | R | 35 | -74 | -6 | 26.473* |
| | R | 34 | -78 | 21 | 45.032** |
| Middle frontal gyrus | L | -43 | 6 | 20 | 30.045* |
| | R | 44 | 5 | 36 | 34.268* |
| | L | -45 | 2 | 52 | 35.226** |
| Postcentral sulcus | L | -30 | -43 | 42 | 52.109*** |
| Superior/Middle frontal gyrus | R | 7 | 8 | 50 | 28.256* |
| Superior parietal lobule | L | -14 | -63 | 43 | 31.112* |

* p<.0001
** p<.00001
*** p<.000001

**Table S2. Three-way repeated measure ANOVA on accuracies**. The table shows the main effects and interactions between the 3 factors (task, skin color and emotion) for the accuracies values within participants (between groups df =1, within groups df = 19).

| Source | F(1,19) |
|---|---|
| Task | 40.055*** |
| Skin | 28.884*** |
| Emotion | 14.085* |
| Task * Skin | 35.677*** |
| Task * Emotion | 12.024* |
| Skin * Emotion | 11.596* |
| Task * Skin * Emotion | 16.994* |

$^{*}$ p<.005

$^{**}$ p<.0001

$^{***}$ p<.00001

**Table S3. Three-way repeated measure ANOVA on response times**. The table shows the main effects and interactions between the 3 factors (task, skin color and emotion) for the response times within participants (between groups df =1, within groups df = 19).

| Source | F(1,19) | p |
|---|---|---|
| Task | 34.587 | * |
| Skin | 3.309 | .085 |
| Emotion | 6.760 | .018 |
| Task * Skin | 30.339 | * |
| Task * Emotion | 4.660 | .044 |
| Skin * Emotion | 5.725 | .027 |
| Task * Skin * Emotion | 4.836 | .040 |

$^{*}$ p<.0001

**Table S4. Descriptive statistics of accuracies conditions**. The table shows the different mean accuracies values, their standard deviations and standard error mean for the different conditions, computed across subjects.

| | Mean | Std. Deviation | Std. Error Mean |
|---|---|---|---|
| Explicit | .893 | .156 | .017 |
| Implicit | .986 | .027 | .003 |
| Angry | .910 | .158 | .017 |
| Happy | .969 | .052 | .005 |
| Black | .911 | .155 | .017 |
| White | .968 | .063 | .007 |
| Angry_explicit | .825 | .189 | .030 |
| Happy_explicit | .961 | .065 | .010 |
| Angry_implicit | .996 | .013 | .002 |
| Happy_implicit | .977 | .033 | .005 |
| Black_explicit | .830 | .186 | .029 |
| White_explicit | .956 | .082 | .013 |
| Black_implicit | .992 | .018 | .002 |
| White_implicit | .981 | .032 | .005 |

**Table S5. Results of different accuracies contrasts**. The table shows the statistics of the difference between mean accuracies for different contrasts, performed with a pair sample t-test computed across subjects.

| Contrasts | t | df | Sig. (2-tailed) |
|---|---|---|---|
| Explicit - Implicit | -5.050 | 79 | * |

| | | | |
|---|---|---|---|
| Angry - Happy | -3.243 | 79 | .002 |
| Black - White | -2.904 | 79 | .005 |
| Angry_explicit - Happy_explicit | -4.333 | 39 | * |
| Angry_explicit - Angry_implicit | -5.562 | 39 | * |
| Happy_explicit - Happy_implicit | -1.373 | 39 | .178 |
| Angry_implicit - Happy_implicit | 3.402 | 39 | .002 |
| Black_explicit - White_explicit | -3.483 | 39 | .001 |
| Black_implicit - White_implicit | 1.746 | 39 | .089 |
| Black_explicit - Black_implicit | -5.350 | 39 | * |
| White_explicit - White_implicit | -1.645 | 39 | .108 |

* p<.0001

**Table S6. Descriptive statistics of response times conditions**. The table shows the different mean response times values, their standard deviations and standard error mean for the different conditions, computed across subjects.

| | Mean | Std. Deviation | Std. Error Mean |
|---|---|---|---|
| Explicit | 843.012 | 111.779 | 12.497 |
| Implicit | 717.357 | 85.444 | 9.552 |
| Angry | 796.617 | 130.255 | 14.563 |
| Happy | 763.753 | 101.373 | 11.333 |
| Black | 790.673 | 126.178 | 14.107 |
| White | 769.696 | 107.917 | 12.065 |
| Angry_explicit | 873.653 | 114.804 | 18.152 |
| Happy_explicit | 812.372 | 101.012 | 15.971 |

| | | | |
|---|---|---|---|
| Angry_implicit | 719.581 | 94.944 | 15.012 |
| Happy_implicit | 715.134 | 75.921 | 12.004 |
| Black_explicit | 875.300 | 102.180 | 16.156 |
| White_explicit | 810.724 | 112.828 | 17.839 |
| Black_implicit | 706.047 | 84.371 | 13.340 |
| White_implicit | 728.668 | 86.068 | 13.608 |

**Table S7. Results of different response times contrasts**. The table shows the statistics of the difference between mean response times for different contrasts, performed with a pair sample t-test computed across subjects.

| **Contrasts** | **t** | **df** | **Sig. (2-tailed)** |
|---|---|---|---|
| Explicit - Implicit | 8.632 | 79 | * |
| Angry - Happy | 2.948 | 79 | .004 |
| Black - White | 2.046 | 79 | .044 |
| Angry_explicit - Happy_explicit | 3.232 | 39 | .003 |
| Angry_explicit - Angry_implicit | 6.427 | 39 | * |
| Happy_explicit - Happy_implicit | 6.245 | 39 | * |
| Angry_implicit - Happy_implicit | .439 | 39 | .663 |
| Black_explicit - White_explicit | 4.256 | 39 | * |
| Black_explicit - Black_implicit | 8.595 | 39 | * |
| White_explicit - White_implicit | 4.247 | 39 | * |
| Black_implicit - White_implicit | -2.285 | 39 | .028 |

* $p<.0001$

**Table S8. Regions of interest peak voxel across folds.** For each of the 5 folds and for each ROI we report peak voxel's coordinates of the classification accuracies produced by the GNB of Explicit vs. Implicit conditions tested against chance level, at q(FDR) < .01 The degrees of freedom were 15 and p-values were all less than .001.

| **Brain Regions** | **L/R** | **x** | **y** | **z** | **t(15)** |
|---|---|---|---|---|---|
| **Extrastriate body area** | | | | | |
| CV1 | R | 58 | -53 | -2 | 6.928** |
| | L | -44 | -66 | 1 | 7.850*** |
| CV2 | R | 56 | -58 | -5 | 6.756** |
| | L | -44 | -67 | 1 | 11.562*** |
| CV3 | R | 59 | -50 | -6 | 5.989* |
| | L | -44 | -66 | 1 | 7.647*** |
| CV4 | R | 48 | -67 | -4 | 6.915** |
| | L | -53 | -62 | -5 | 9.740*** |
| CV5 | R | 53 | -60 | -4 | 7.531** |
| | L | -45 | -65 | 1 | 8.781*** |
| **Ventrolateral prefrontal cortex** | | | | | |
| CV1 | R | 46 | 13 | 27 | 9.289*** |
| CV2 | R | 48 | 14 | 26 | 8.919*** |
| CV3 | R | 48 | 14 | 26 | 8.250*** |
| CV4 | R | 48 | 15 | 26 | 11.163*** |
| CV5 | R | 50 | 12 | 26 | 9.831*** |

| | | | | | |
|---|---|---|---|---|---|
| **Intraparietal sulcus** | | | | | |
| CV1 | R | 35 | -72 | 37 | 6.419* |
| CV2 | R | 35 | -73 | 36 | 6.267* |
| CV3 | R | 34 | -72 | 37 | 8.918*** |
| CV4 | R | 35 | -73 | 36 | 6.623** |
| CV5 | R | 35 | -75 | 35 | 9.503*** |
| **Precuneus** | | | | | |
| CV1 | R | 5 | -67 | 49 | 7.254** |
| CV2 | R | 6 | -67 | 49 | 6.359* |
| CV3 | R | 6 | -67 | 48 | 5.846* |
| CV4 | R | 8 | -68 | 49 | 7.225** |
| CV5 | R | 7 | -69 | 52 | 6.993** |
| **Inferior parietal lobule** | | | | | |
| CV1 | R | 53 | -49 | 24 | 6.458* |
| CV2 | R | 53 | -49 | 24 | 6.644** |
| CV3 | R | 43 | -49 | 38 | 6.465** |
| CV4 | R | 50 | -49 | 26 | 7.254** |
| CV5 | R | 54 | -49 | 26 | 7.793*** |

* p<.0001
** p<.00001
*** p<.000001

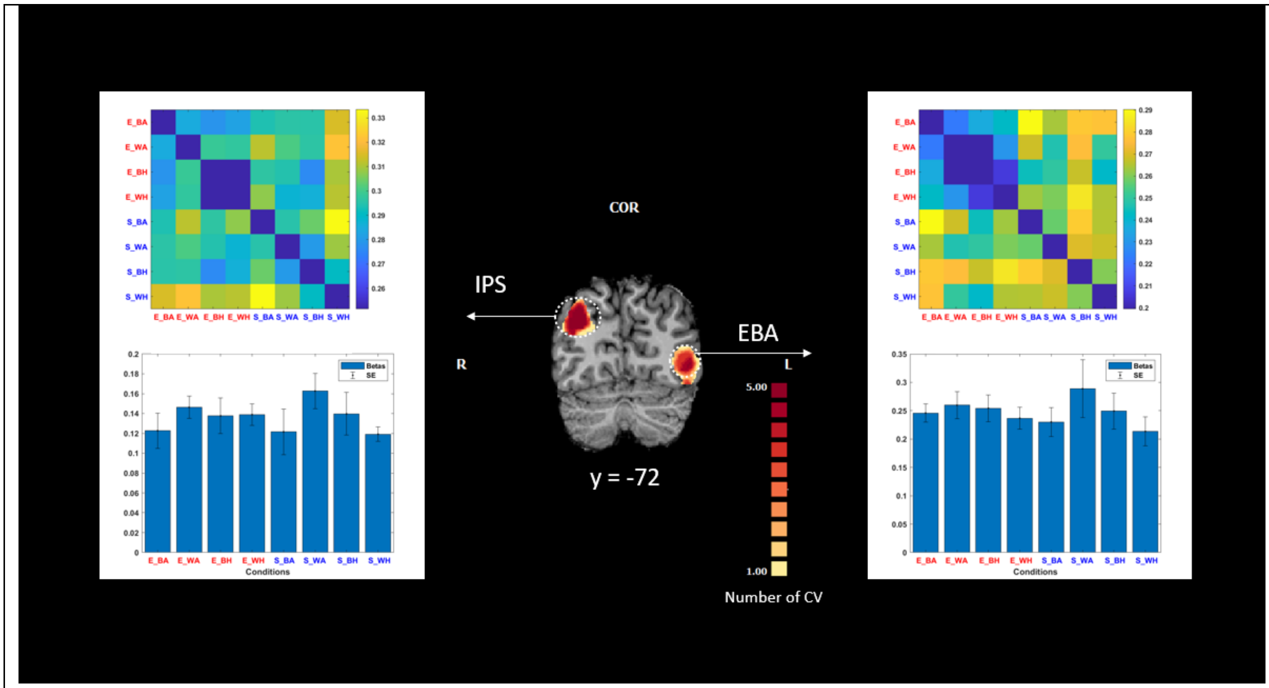


**Figure S1. Details of the responses from the ROIs identified by the decoding of task effect. RDM and beta plots at the category level of each ROIs are shown.** Results shown in these images were obtained by following the same procedures as described in the main text (see Fig. 8). The RDM for (left) EBA shows a pattern of similarities within the explicit condition but the differences in the beta values between the tasks are not significant. Abbreviations: EBA = extrastriate body area; IPL = inferior parietal lobe; IPS = intraparietal sulcus.

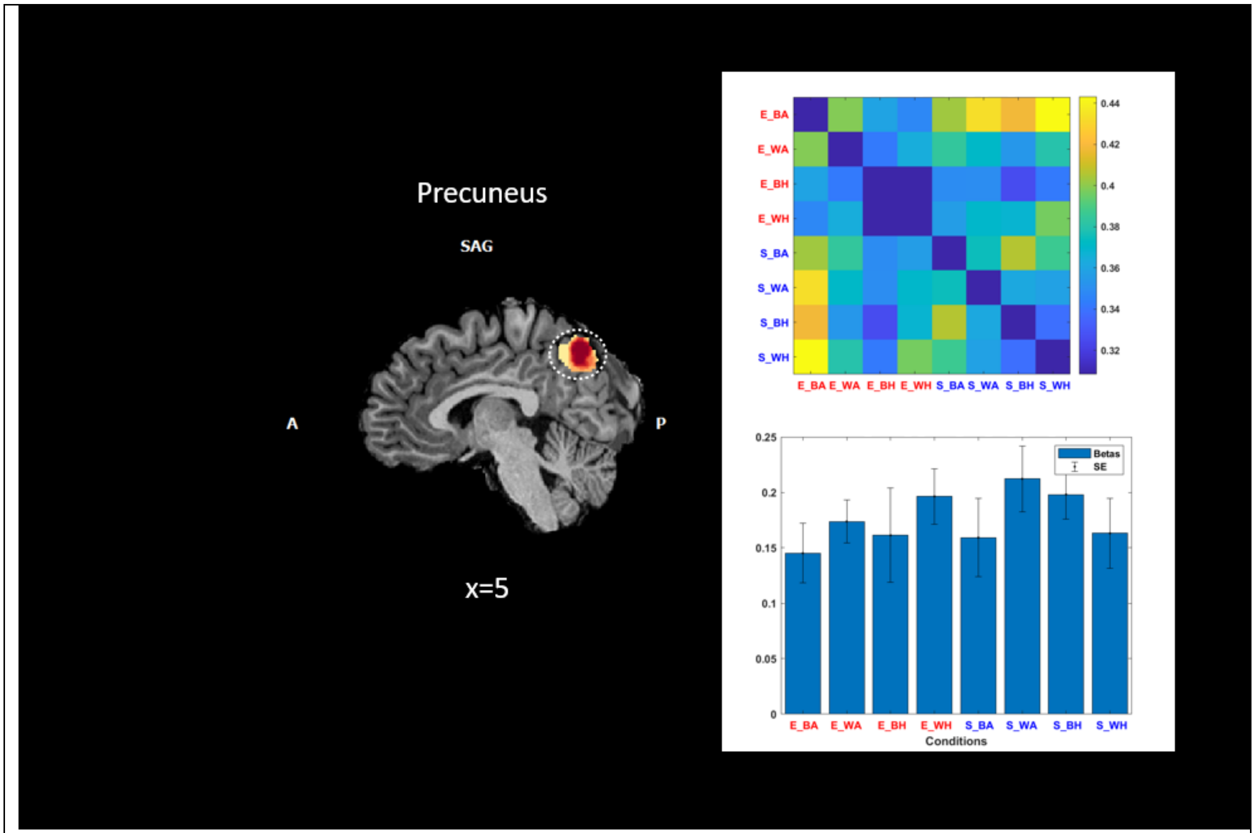


**Figure S2. Details of the responses from the ROIs identified by the decoding of task effect, RDM and beta plots at the category level of each ROIs is shown.** For the precuneus we did not find a similarity pattern and differences in beta plots are not significant.
